# RNAi screen of RING/U-box domain ubiquitin ligases identifies critical regulators of tissue regeneration in planarians

**DOI:** 10.1101/2021.10.12.463740

**Authors:** John M. Allen, Madison Balagtas, Elizabeth Barajas, Carolina Cano Macip, Sarai Alvarez Zepeda, Ionit Iberkleid, Elizabeth M. Duncan, Ricardo M. Zayas

**Author notes:** **Correspondence**: Dr. Ricardo M. Zayas.

## Abstract

Regenerative processes depend on the interpretation of signals to coordinate cell behaviors. The role of ubiquitin-mediated signaling is known to be important in many cellular and biological contexts, but its role in regeneration is not well understood. To investigate how ubiquitylation impacts tissue regeneration *in vivo*, we are studying planarians that are capable of regenerating after nearly any injury using a population of stem cells. Here we used RNAi to screen RING/U-box E3 ubiquitin ligases that are highly expressed in planarian stem cells and stem cell progeny. RNAi screening identified nine genes with functions in regeneration, including the spliceosomal factor *prpf19* and histone modifier *rnf2*; based on their known roles in developmental processes, we further investigated these two genes. We found that *prpf19* was required for animal survival but not for stem cell maintenance, suggesting a role in promoting cell differentiation. Because RNF2 is the catalytic subunit of the Polycomb Repressive Complex 1 (PRC1), we also examined other putative members of this complex (CBX and PHC). We observed a striking phenotype of regional tissue misspecification in *cbx* and *phc* RNAi planarians. To identify genes regulated by PRC1, we performed RNA-seq after knocking down *rnf2* or *phc*. Although these proteins are predicted to function in the same complex, we found that the set of genes differentially expressed in *rnf2* versus *phc* RNAi were largely non-overlapping. Using *in situ* hybridization, we showed that *rnf2* regulates gene expression levels within a tissue type, whereas *phc* is necessary for the spatial restriction of gene expression, findings consistent with their respective *in vivo* phenotypes. This work not only uncovered roles for RING/U-box E3 ligases in stem cell regulation and regeneration, but also identified differential gene targets for two putative PRC1 factors required for maintaining cell-type-specific gene expression in planarians.

## 1 Introduction

A deep understanding of the networks and signaling pathways that direct the maintenance and differentiation of adult stem cells is essential for regenerative therapies. The freshwater planarian, *Schmidtea mediterranea*, is an important model for studying the molecular mechanisms that underpin stem cell-based regeneration (Elliott and Sánchez Alvarado, 2013; Ivankovic et al., 2019). These worms maintain a large population of adult stem cells, a subset of which have been demonstrated to be pluripotent (Baguñà et al., 1989; Wagner et al., 2011). This population of stem cells continuously renews planarian tissues during homeostasis and is also mobilized in response to injury to regenerate tissues (Saló and Baguñà, 1985; Abnave et al., 2017). As such, they offer an amenable model to study stem cell biology in a whole-organism *in vivo* context.

Extensive work has been performed to understand the molecular basis of planarian regeneration (Reddien, 2018), yet most studies have primarily examined transcriptional changes (Labbé et al., 2012; Onal et al., 2012; van Wolfswinkel et al., 2014; Fincher et al., 2018; Plass et al., 2018). Comparatively, fewer studies have focused on proteomic regulation in planarian stem cells (Fernandez-Taboada et al., 2011; Boser et al., 2013) or the post-translational regulation of proteins vital for stem cell function (Strand et al., 2018). One essential post-translation regulator of proteins is the addition of the small, highly conserved polypeptide ubiquitin, which modifies protein function in myriad cellular contexts, including transcription, cell cycle regulation, translational fidelity, protein turnover, and degradation (Ciechanover et al., 1984; Nakayama and Nakayama, 2005; Endoh et al., 2012; Higgins et al., 2015).

Ubiquitin-dependent signaling events have emerged as essential regulators of stem cell functions, including self-renewal and differentiation (Werner et al., 2017). The transfer of free ubiquitin onto a target substrate typically occurs through a tripartite enzymatic cascade that terminates with the E3 ubiquitin ligases. The E3 ligases can be grouped into two major classes: the HECT (Homologous to the E6-AP Carboxyl Terminus) and the more prevalent RING (Really Interesting New Gene) class. Of the approximately 617 genes encoding putative E3 ligases identified in the human genome, 309 were predicted to contain a RING finger (RNF) or the related U-box domain; a further 270 E3 genes that function in complexes associated with RINGs (Li et al., 2008). The RNFs are defined by a zinc-finger domain with an evolutionarily conserved arrangement of cysteine and histidine residues that coordinate two zinc ions and bind an E2-ubiquitin conjugate (Lorick et al., 1999). The U-box domain forms a similar structure to the RING domain and can bind conjugated E2 but does not coordinate zinc (Aravind and Koonin, 2000). Substrate recognition and binding are achieved by additional domains within the RNF protein or association with other proteins as part of a multi-protein complex. Previous work on E3 ligase function in planarians has implicated a subset of HECT E3 and Cullin-RING complex member ligases as essential regulators of regeneration and stem cells (Henderson et al., 2015; Strand et al., 2018).

Here we performed functional analysis on a subgroup of RING and U-box domain-containing genes expressed in the planarian stem cells or progeny. We found several to be essential for homeostatic maintenance, regeneration, and tissue patterning, including spliceosomal factor *prpf19* and epigenetic factors *rnf2* and *bre1*, known to ubiquitylate histones H2A and H2B, respectively. *prpf19* was required for worm survival but not for stem cell maintenance, suggesting a role in promoting cell differentiation. In addition, the Polycomb Repressive Complex 1 (PRC1) gene *rnf2* was required for global monoubiquitylation of histone H2A (H2Aub1) and promoting proper regeneration. In contrast, when we disrupted putative PRC1 genes *phc* and *cbx*, we did not detect a global reduction in H2Aub1 levels but did observe specific defects in the organization of tissue near the base of the planarian pharynx. Taken together, analysis of RING/U-box E3 ligases identified multiple regulators of stem cell biology and regeneration and led to the discovery of differential phenotypes and transcriptional targets for putative PRC1 factors.

## 2 Materials and Methods

### 2.1 Planarian care

A clonal line of asexual *S. mediterranea* (CIW4) was used in all experiments and kept in 1X Montjuïc salts (1.6 mM NaCl, 1.0 mM CaCl_2_, 1.0 mM MgSO_4_, 0.1 mM MgCl_2_, 0.1 mM KCl, 1.2 mM NaHCO_3_, pH 7.0) (Cebrià and Newmark, 2005) in food-grade plastic containers at 20°C (Merryman et al., 2018). Animals selected for experiments were 3-6 mm in length and starved for one week before experimentation.

### 2.2 Gene identification and cloning

To find RING and U-box domain-containing genes in *S. mediterranea*, we filtered the Dresden transcriptome (Brandl et al., 2016; Rozanski et al., 2019) using InterPro Domain IDs (Blum et al., 2021), IPR001841 (Zinc finger, RING-type), and IPR003613 (U box domain). This list was filtered to include only the longest gene contig for each hit and was used as query sequences for a BLAST search to a curated list of human RING and U-box genes (Li et al., 2008) at an expected value cut-off of 1 × 10^−3^. We additionally filtered the Dresden transcriptome for contigs annotated with IPR013083 (Zinc finger, RING/FYVE/PHD-type). This list was filtered to remove duplicate entries, and a BLAST search was performed against our list of human RING and U-box genes as the IPR013083 family contains non-RING and U-box genes, only genes that had predicted homology to a human gene at an expected cut-off of 1 × 10^−3^ were appended to our initial list (Supplementary Table S1).

The sequences of interest were obtained from either an EST library (Zayas et al., 2005) or cloned using gene-specific primers into pPR-T4P using ligation-independent cloning (Liu et al., 2013; Adler and Sánchez Alvarado, 2018). EST clone accession numbers and the primer sequences used are listed in Supplementary Tables S2-S3.

### 2.3 RNA interference

During the initial screening, animals were fed double-stranded RNA (dsRNA) mixed with a ≈3:1 mixture of liver-water paste twice per week for eight feeds and were amputated pre-pharyngeally on day 28 of treatment to observe regeneration. *In vitro* transcribed and dsRNA expressed in bacteria were used to perform RNAi during the initial screening of RING and U-box genes; all subsequent RNAi knockdowns were performed using dsRNA expressed in bacteria. *In vitro* dsRNA was synthesized as previously described (Rouhana et al., 2013); the entire reaction mixture was separated into eight aliquots, mixed with liver paste, and stored until feeding. Bacterially-expressed dsRNA was prepared by growing *E. coli* strain HT115 transformed with the pPR-T4P plasmid (Liu et al., 2013; Adler and Sánchez Alvarado, 2018) containing the gene of interest and inducing dsRNA expression using IPTG. Bacteria pellets were purified using centrifugation and mixed with liver paste for administration to animals (Gurley et al., 2008).

### 2.4 In situ hybridization

Antisense probes for *in situ* hybridization were synthesized as previously described (Pearson et al., 2009) from DNA templates amplified from pBS II SK(+) (Stratagene) or pPR-T4P (Liu et al., 2013; Adler and Sánchez Alvarado, 2018) plasmid vectors incorporating either digoxigenin- or FITC-labeled UTPs. Animals for whole-mount in situ hybridization (WISH) were processed and hybridized as outlined previously (King and Newmark, 2013). Briefly, samples were sacrificed in 5% *n*-acetyl cysteine in 1X PBS, fixed in 4% formaldehyde in PBS with 0.3% Triton X-100 (PBS-Tx), and bleached in a formamide/hydrogen peroxide bleaching solution (5% deionized formamide, 1.2% H_2_O_2_, in 0.5X SSC). Samples were pre-hybridized for two hours and then hybridized with probe overnight at 56°C. Next, samples were incubated with an appropriate antibody, depending on the probe label and subsequent development strategy. For chromogenic development, samples were incubated with an anti-digoxigenin-AP antibody (Roche, 1:2000) and developed with NBT/BCIP in AP buffer. Fluorescent *in situ* development was performed using Fast Blue (Lauter et al., 2011) or Tyramide Signal Amplification (TSA) after incubation with anti-digoxigenin-AP or anti-FITC-POD (Roche, 1:300) antibodies, respectively, following previously described protocols(King and Newmark, 2013; Brown and Pearson, 2015). For irradiation experiments to eliminate dividing cells, worms were exposed to 60 Gy of X-ray irradiation in a Precision CellRad Irradiation System and processed for WISH 7 days post-irradiation.

### 2.5 Anti-phosphohistone H3 immunohistochemistry

Animals were incubated in ice-cold 2% hydrochloric acid for 5 minutes and fixed for 2 hours in Carnoy’s solution (60% ethanol, 30% chloroform, 10% glacial acetic acid), at 4°C. Samples were washed in methanol for 1 hour at 4°C and bleached overnight in 6% H_2_O_2_ diluted in methanol at room temperature. Animals were washed out of methanol and into PBS-Tx and blocked in 1% bovine serum albumin (BSA) diluted in PBS-Tx for 4 hours at room temperature. Samples were incubated with anti-phosphohistone H3 (Ser 10) (Cell Signaling 3377, 1:1000) diluted in 1% BSA/PBS-Tx overnight at 4°C. Washes were performed using PBS-Tx (6 × 1 hour), and Samples were washed extensively in PBS-Tx (6 × 1 hour) and incubated with anti-rabbit-HRP (Cell Signaling #7074, 1:1000) diluted in 1% BSA/PBS-Tx. Signal was developed using TSA as previously described (King and Newmark, 2013).

### 2.6 TUNEL staining

The Terminal Deoxynucleotidyl Transferase-mediated deoxyuridine triphosphate Nick End-labeling (TUNEL) assay was performed to quantify apoptotic cells. Animals were incubated in 5% *n*-acetyl cysteine in PBS for 5 minutes and fixed in 4% formaldehyde diluted in PBS-Tx for 15 minutes.

Samples were then permeabilized in 1% SDS diluted in PBS and bleached overnight in 6% H_2_O_2_ in PBS-Tx. As previously described, samples were then rinsed and stained using the ApopTag Kit (Millipore-Sigma) (Pellettieri et al., 2010).

### 2.7 Protein extraction and western blotting

RNAi planarians were homogenized in TRIzol (ThermoFisher). The organic phase was recovered following the manufacturer-provided TRIzol protocol with a modified solubilization buffer (4M Urea, 0.5% SDS) to isolate proteins for western blot. An added sonication step of 10 one-second pulses was performed to increase protein recovery (Simoes et al., 2013; Duncan et al., 2015). Samples were loaded onto AnyKD TGX gels (BioRad), transferred using the semidry method to a 0.45m PVDF membrane, and blocked in 5% nonfat milk/TBS-Tw (Tris-buffered saline with 0.1% Tween-20). Antibodies to monoubiquityl-Histone H2A (Cell Signaling 8240), monoubiquityl-Histone H2B (Cell Signaling 5546), and anti-Ubiquitin (Cell Signaling #3933) were diluted in 5% bovine serum albumin in TBS-Tw at 1:2000, 1:1000 and 1:1000 respectively and incubated overnight at 4°C. Washes were performed with TBS-Tw and anti-rabbit-HRP (Cell Signaling #7074) was diluted in 5% nonfat milk/TBS-Tw at 1:2500 and incubated for 1 hour at room temperature. Signal was developed using BioRad Clarity Western ECL Substrate (BioRad 1705061). Loading was normalized to total protein for monoubiquityl-Histone H2A and Ubiquitin blots using AnyKD TGC Stain-Free gels (BioRad). Loading for monoubiquityl-Histone H2B blots was normalized to mouse anti-β-tubulin (1:1000 dilution, DSHB #E7) with anti-mouse-HRP secondary (1:1000 dilution, Cell Signaling #7076).

### 2.8 RNA sequencing

Worms from three independent control and experimental RNAi groups per time point were homogenized in TRIzol, and RNA was extracted and purified following manufacturer protocol. RNA was treated with the Turbo DNA-free kit and column purified using the Qiagen RNeasy MinElute Cleanup kit. Three independent biological groups were collected at each time point assayed for both control and experimental (*rnf2* or *phc*) RNAi treatments. Samples were sequenced on an Illumina HiSeq 4000 to a read depth of at least 15 million 150 bp paired-end reads. The sequenced reads were submitted to the NCBI BioProject PRJNA768725. Reads were pseudoaligned to the Dresden (dd_Smed_v6) transcriptome using kallisto (Bray et al., 2016), and differential gene expression analysis was performed using the R Bioconductor package (Huber et al., 2015) and DESeq2 (Love et al., 2014) with an FDR cut-off value of ≤ 0.1 applied. To perform Gene Ontology (GO) analysis, differentially expressed transcripts from the day 28 *rnf2(RNAi)* data set were compared to the human proteome using BLASTX (cut-off e-value < 1e^-3^). Human UniProt IDs were used as input for annotation and overrepresentation analysis (http://geneontology.org/) using Fisher’s Exact test with an FDR multiple comparisons correction cut-off of ≤ 0.05 applied.

### 2.9 Reverse transcription quantitative PCR

Total RNA was extracted and purified from whole worms as described above. cDNA was synthesized using the iScript Reverse Transcription Supermix for RT-qPCR Kit (BioRad #1708840). Reverse transcription quantitative PCR (RT-qPCR) was performed on a Bio-Rad CFX384 Touch Real-Time PCR Detection System using iTaq Universal SYBR Green Supermix (BioRad #1725120) with two-step cycling protocol with an annealing/extension temperature of 60.0°C. Three biological and three technical replicates were performed for each experiment. The relative amount of each target was normalized to *β-tubulin* (accession # DN305397), and normalized relative expression changes were calculated using the ΔΔCq method (Livak and Schmittgen, 2001). Significance was determined at a p-value < 0.05 using Student’s t-test with Holm-Sidak correction for multiple comparisons.

## 3 Results

### 3.1 Identification of RING and U-Box E3 ubiquitin ligase genes in *S. mediterranea*

The RING and U-Box protein domains have been identified as having a pivotal role in mediating the ubiquitylation of a target substrate (Lorick et al., 1999; David et al., 2011). To identify genes in *S. mediterranea* that are predicted to encode a RING/U-box domain, we filtered a reference planarian transcriptome (Brandl et al., 2016) using InterPro domain annotations and generated a list of 393 transcripts. Next, we used the predicted RING and U-box domain-containing gene transcripts to perform BLAST analysis against a curated list of human E3 ubiquitin ligases (Li et al., 2008). We found 376 planarian genes that were predicted to have homology with a human RING/U-box gene (Supplementary Table S1) and 17 planarian transcripts that, while having predicted RING or U-box domains, did not have predicted significant homology to a human RING/U-box gene. Finally, we classified these putative planarian RINGs into major subfamilies based on their homology to human genes and found representative factors for most (15/17) subfamilies (Supplementary Table S1).

### 3.2 A functional screen reveals genes with roles in planarian stem cell regulation and regeneration

To identify RING/U-box genes that regulate planarian stem cell function in tissue maintenance and regeneration, we assessed the function of 93 genes from our list (≈25%). To better identify factors that regulate regeneration, we included 72 genes predicted to be expressed in stem cells and stem cell progeny (Supplementary Table S2) based on data from a sorted-cell transcriptome (Labbé et al., 2012). RNAi treatments were performed over four weeks while the worms were monitored for defects in homeostasis. After 28 days, planarians were amputated to assess the effect of RNAi treatment on regeneration (Figure 1A). We found that RNAi of nine genes produced phenotypes related to stem cell function in homeostasis during regeneration (Table 1). Phenotypes observed during homeostasis included head regression, epidermal lesions, ventral curling, and lysis (Figure 1B); other genes displayed abnormalities and delays during regeneration when disrupted (Figure 1C). During homeostasis, head regression was observed after RNAi-mediated targeting of *prpf19, march5, traf-2A/B, not4, rnf8-like*, and *bre1*; lesions were observed after disruption of *march5, ran*, and *bre-1*; and ventral curling was observed after disruption of *prpf19* and *not4*. The genes *prpf19, march5*, and *ran* were essential for worm survival, and depletion of these transcripts caused worm lysis.

**Figure 1.**
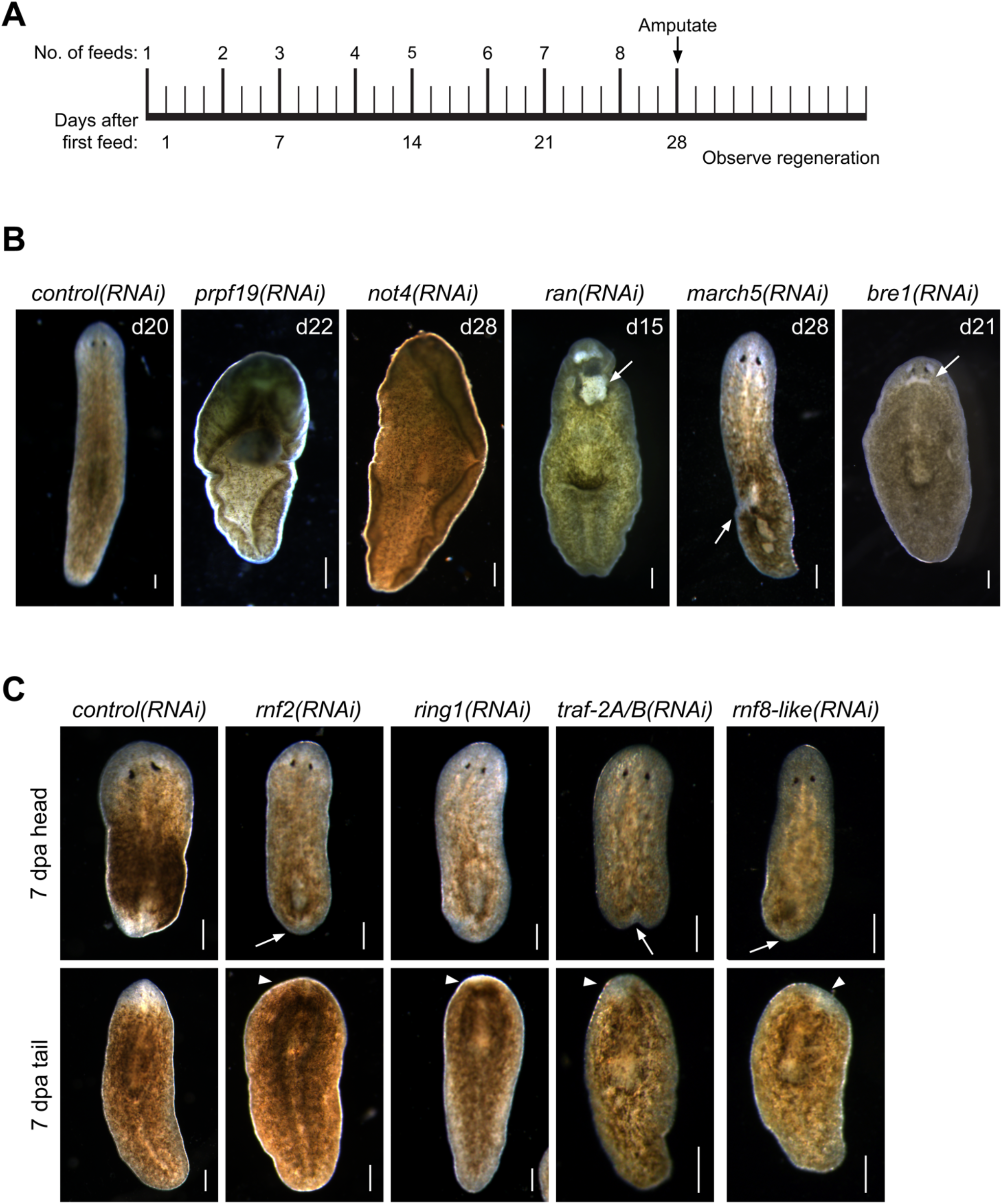
RNAi screen of RING/U-Box E3 ubiquitin ligases identifies regulators of stem cells and regeneration. (A) Feeding and amputation schedule of RNAi screen. Worms were fed twice per week for a total of eight feeds and amputated pre-pharyngeally on day 28. (B) Knockdown of the indicated genes resulted in phenotypes, including ventral curling (N = 33/121 and 7/37 for *prpf19* and *not4*, respectively) and lesions (white arrow, N = 11/43, 18/33, and 14/53 for *ran, march5*, and *bre1*, respectively). Animals are shown after the conclusion of the RNAi feedings and before amputation. (C) Knockdown of the indicated genes that demonstrated phenotypes of delayed or absent regeneration after amputation, as shown by the smaller than normal or absent blastemas (white arrow) and missing or faint eyespots (white arrowhead) when compared to *control(RNAi)* worm at the same regeneration time point (N = 37/58, 19/29, 31/36, and 21/30 trunk fragments for *rnf2, ring1, traf-2A/B*, and *rnf8-like*, respectively). Scale bars = 200 μm.

**Table 1:** RING/U-box E3 ubiquitin ligases showing phenotypes following RNAi.

| Gene Name | Dresden Contig ID | Human RING or U-box Homolog | E-value | Phenotypes Observed |
| --- | --- | --- | --- | --- |
| *Smed-bre1* | dd_Smed_v6_4070_0_1 | RNF40 | 1e-88 | HR, DR |
| *Smed-march5* | dd_Smed_v6_4602_0_1 | MARCH5 | 8e-93 | HR, Lesions, Lysis |
| *Smed-not4* | dd_Smed_v6_4767_0_1 | CNOT4 | 5e-87 | HR, VC |
| *Smed-prpf19* | dd_Smed_v6_1276_0_1 | PRPF19 | 0.0 | HR, VC, Lysis |
| *Smed-ran* | dd_Smed_v6_330_0_1 | CBLB* | 3e-94 | Lesions, Lysis |
| *Smed-ring1* | dd_Smed_v6_12141_0_1 | RING1 | 6e-33 | DR |
| *Smed-rnf2* | dd_Smed_v6_8989_0_1 | RNF2 | 6e-46 | DR |
| *Smed-rnf8-like* | dd_Smed_v6_1137_0_5 | RNF8 | 4e-05 | DR, Lesions |
| *Smed-traf-2A/B* | dd_Smed_v6_3837_0_1 | TRAF2 | 2e-69 | HR, DR |

HR: Head regression. VC: Ventral Curling. DR: Delayed regeneration; \*Top Human BLAST hit: RAN, e-value: 9e-116

Knockdown of *rnf8-like, bre1, rnf2*, and *ring1* caused defective regeneration, typically manifested as a delayed appearance of visible eyespots compared to *control(RNAi)* treatments. The genes that demonstrated phenotypes (Table 1) were then examined by WISH. All had broad expression patterns but showed discrete expression in major differentiated tissue types like the cephalic ganglia or the intestine (Figure 2A). We chose to analyze further the *prpf19* phenotype as it was predicted to be expressed in stem cells and its phenotype of ventral curling suggested a role in regulating stem cells, and the *bre1* and *rnf2* phenotypes based on their known roles as epigenetic regulators during developmental processes.

**Figure 2.**
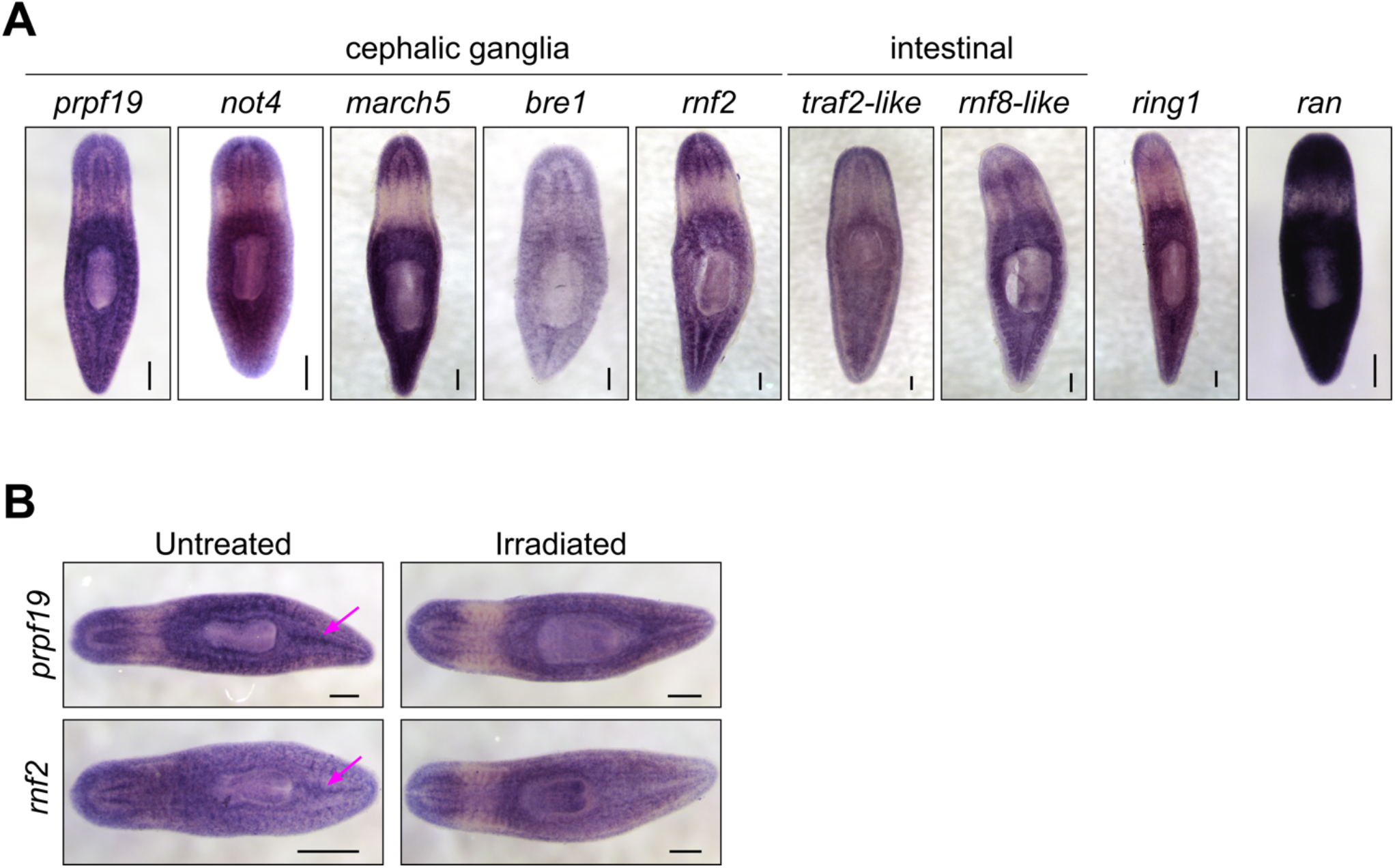
(A) WISH expression patterns for genes showing phenotypes in the RNAi screen. All the genes examined were expressed throughout the parenchyma; a subset of genes displayed enriched expression near the cephalic ganglia or the intestine. (B) WISH analysis of *prpf19* and *rnf2* in untreated controls (right) and irradiated worms (left). Arrows show expression in regions enriched in stem cells in untreated worms that are undetectable in irradiated worms. Scale bars = 200 μm.

### 3.3 Spliceosomal factor *prpf19* is required for worm survival and stem cell function

The U-box gene *prpf19* had enriched expression in planarian stem cells, and our initial screen revealed that its expression was required for worm survival. Other aspects of the RNAi phenotype, including head regression and ventral curling, are typically associated with the loss of stem cells or their function. These phenotypes are consistent with an earlier report for *prpf19* as being up-regulated during and necessary for head regeneration in planarians (Roberts-Galbraith et al., 2016). In other organisms, *prpf19* encodes a core component of the NineTeen Complex (NTC), with a well-described role in regulating mRNA splicing. Consistent with a role in an essential cellular process, we found broad expression of this gene using WISH (Figure 2A).

Consistent with our analysis of a published sorted-cell transcriptome (Labbé et al., 2012), we found that at least a subset of this expression is in the stem cells or stem cell progeny by performing WISH on worms seven days after irradiation treatment (Figure 2B). We confirmed this observation using double fluorescent in situ hybridization (FISH) to observe co-expression of *prpf19* with stem cell markers *piwi-1* and *h2b*, and stem cell progeny markers *prog-1* and *agat-*1 (Supplementary Figure S2A-B). As *prpf19* has been shown to function as an E3 ubiquitin ligase (Song et al., 2010), we assayed the effect of *prpf19* RNAi on ubiquitylated proteins in whole-worm protein extracts by western blotting using a pan-ubiquitin antibody. We did not detect changes in ubiquitylation levels compared to controls, suggesting that *prpf19* disruption does not appreciably affect global ubiquitylation or has only a minor effect that is not resolvable on a total ubiquitin blot (Supplementary Figure S2C).

To investigate if the *prpf19(RNAi)* phenotypes observed resulted from stem cell depletion, we performed WISH to stem cell marker genes *tgs-1, piwi-1*, and *h2b* on *prpf19(RNAi)* and control worms. Surprisingly, all marker genes analyzed showed robust expression, even in worms where the phenotype had significantly progressed (Figure 3A). Furthermore, because *prpf19* was found to be expressed in additional cell types besides stem cells (Figure 2), we examined the effect of *prpf19* inhibition on epidermal differentiation by performing WISH with markers for early and late epidermal progeny, *prog-1* and *agat-1*, respectively (Eisenhoffer et al., 2008; van Wolfswinkel et al., 2014). Consistent with the epidermal lesions observed during the progression of the *prpf19* phenotype, staining for epidermal lineage markers was reduced in *prpf19(RNAi)* worms (Figure 3A). In addition, we analyzed relative mRNA levels after *prpf19* RNAi using RT-qPCR for marker genes in the epidermal lineage. We measured the expression of *zfp-*1, which marks epidermal stem cells, progenitor markers *prog-1* and *agat-1*, and the mature epidermal cell marker gene *vim-1*(van Wolfswinkel et al., 2014; Tu et al., 2015). We found that levels of epidermal marker genes were reduced after *prpf19(RNAi)* (Supplementary Figure S1D). Importantly, we did not observe a reduction in the relative expression level of epidermal stem cell marker, *zfp-1*, suggesting *prpf19* inhibition is not causing an appreciable loss of this subset of stem cells. These results indicate that *prpf19* function is not required for the maintenance and survival of planarian stem cells but may affect their differentiation into epidermal progenitors or the maintenance of post-mitotic progenitor populations.

**Figure 3.**
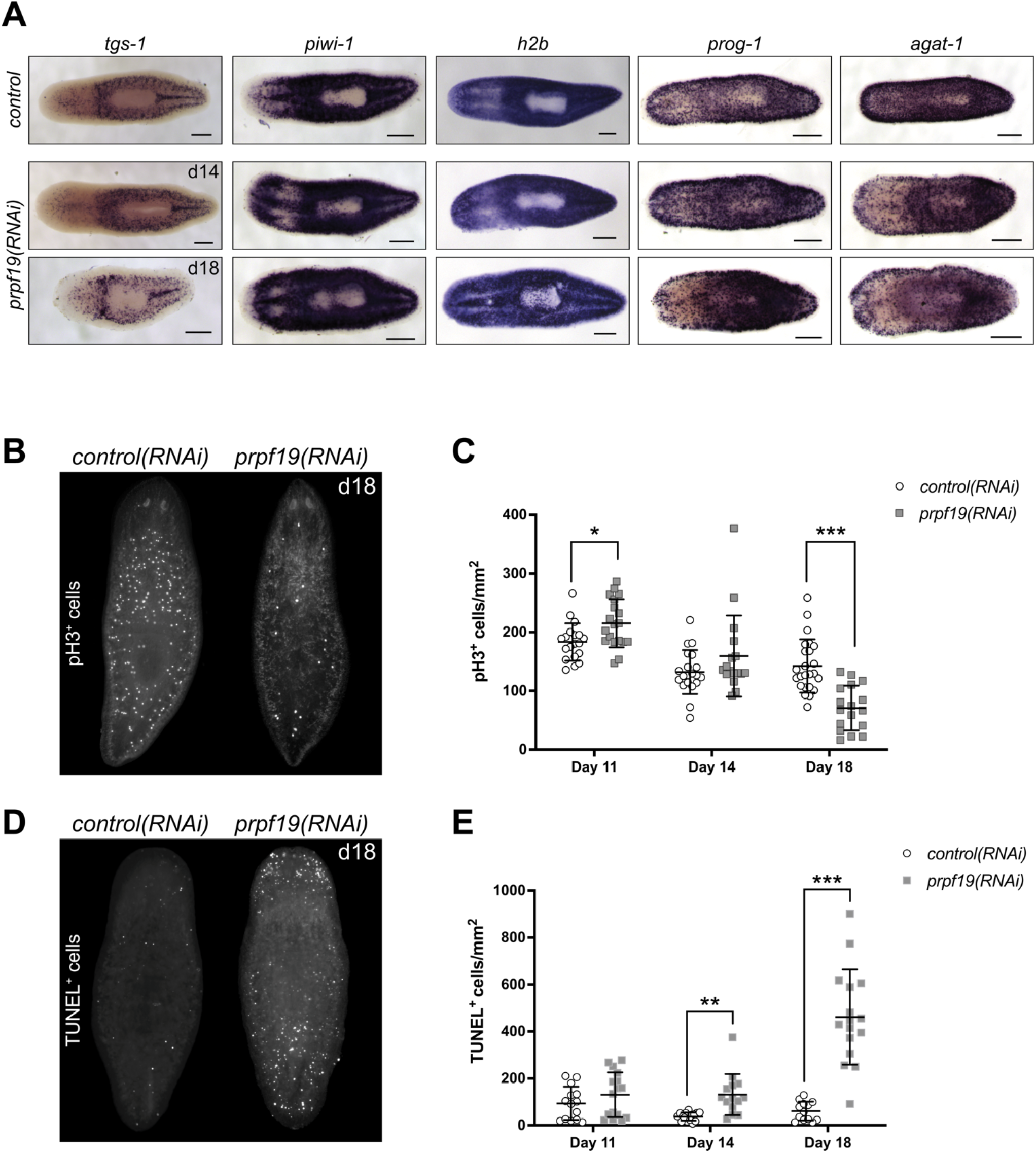
Inhibition of *prpf19* disrupts stem cell function but is not required for stem cell maintenance. (A) WISH to stem cell markers *tgs-1* (N = 10-11), *piwi-1* (N = 7-9) and *h2b* (N = 4), and early and late epidermal progeny markers *prog-1* (N= 7-9) and *agat-1* (N = 8-9), respectively, in *control(RNAi)* (upper panels) and *prpf19(RNAi)* animals at 14 (middle panels) and 18 (bottom panels) days after first RNAi feeding. (B) Representative image of animals fixed 18 days after first RNAi feeding for *control(RNAi)* (left, N = 24) or *prpf19(RNAi)* (right, N = 17) and immunostained for mitotic marker phospho-histone H3. (C) Quantification of phospho-histone H3^+^ cells per mm^2^ of worms fixed at 11, 14, and 18 days after first RNAi feed (N = 17 – 24 per time point). (D) Representative image of animals fixed 18 days after first RNAi feeding for *control(RNAi)* (left, N = 13) or *prpf19(RNAi)* (right, N = 16) and processed for TUNEL staining. (E) Quantification of TUNEL^+^ cells per mm^2^ of worms fixed at 11, 14, and 18 days after first RNAi feed (N = 13 – 16 per time point) All data are represented as mean ± SD. *p-value < 0.05, **p-value < 0.001, ***p-value < 0.0001, Student’s t-test with Holm-Sidak correction for multiple comparisons. Scale bars = 200 μm.

### 3.4 Inhibition of *prpf19* causes defects in stem cell proliferation and an increase in cell death

Despite being dispensable for stem cell maintenance, the strong expression of *prpf19* in stem cells and robust phenotypes that resulted from *prpf19* inhibition suggested a role for *prpf19* in regulating stem cell dynamics. To examine the effect of *prpf19* RNAi on cell proliferation, we stained *control(RNAi)* and *prpf19(RNAi)* worms with anti-phospho-histone H3 (pH3) to mark mitotic cells across several time points days prior to and after the onset of the morphological phenotype. Initially, the animals showed a small but significant increase in pH3^+^ cells (day 11). However, we found that at the later time points (day 18), when the external phenotype is beginning to manifest, there was a significant decrease in the number of pH3^+^ cells in *prpf19(RNAi)* worms (Figure 3B and 3C). Furthermore, this decrease in the number of mitotic cells was not correlated with a reduction in the expression of stem cell marker genes (Figure 3A), suggesting that *prpf19(RNAi)* treatment may block or alter the rate of stem cell differentiation.

To better understand the severe phenotypes observed in *prpf19(RNAi)* worms, including epidermal lesioning and worm lysis, we assayed the worms for dying cells using TUNEL. Not surprisingly, we found an increase in TUNEL^+^ cells in *prpf19(RNAi)* worms compared to control worms at the time point before observing phenotypes, and a marked increase was observed as the *prpf19(RNAi)* phenotype progressed (Figure 3D and 3E). This result is congruous with reports of *prpf19* having anti-apoptotic effects in human cell lines (Lu and Legerski, 2007). Together with the observed loss of epidermal progenitor markers (Figure 3A; Supplementary Figure S1D), the data suggests that the phenotype observed after *prpf19* depletion is not caused by a loss of stem cells. Instead, the observed loss of epidermal integrity may result from abnormal stem cell function, either through impaired differentiation and homeostatic replacement of differentiated tissues or impaired proliferation. In addition, *prpf19* may also have a role as an anti-apoptotic factor in differentiated tissues, resulting in increased apoptosis in *prpf19(RNAi)* worms.

### 3.5 NTC components and targets are necessary for tissue renewal and regeneration

NTC is a large protein complex with various cellular roles but has its best-described role in regulating pre-mRNA splicing. Named after its founding member, *prpf19*, the complex is conserved between humans and yeast. NTC functions as an E3 ligase through its PRPF19 subunit to stabilize the association of snRNP spliceosome components (Figure 4A). To examine if the effects of *prpf19* RNAi were being mediated through disruption of a conserved spliceosomal complex, we knocked down three homologs of core NTC component members, *cdc5l, plrg1*, and *spf27* (Supplementary Table S3). We found that these genes were also necessary for worm survival and regeneration (Figure 4B and 4C). *cdc5l* and *plrg1* are essential for NTC function in yeast and also presented very severe phenotypes in *S. mediterranea*, with RNAi animals phenocopying the head regression, ventral curling, and lysis that we observed after *prpf19* RNAi. *Spf27(RNAi)* worms displayed a milder phenotype than other NTC genes examined and showed delayed or absent regeneration in 28/37 head fragments and 33/37 trunk fragments; an additional three trunk fragments showed a more severe ventral curling and lysis phenotype. Also, we postulated that if the *prpf19* RNAi phenotype resulted from its ubiquityl ligase activity within the NTC complex, inhibiting *prpf3* or *prpf8* should result in a similar phenotype. Indeed, we found that *prpf3(RNAi)* and *prpf8(RNAi)* worms exhibited severe phenotypes like *prpf19* RNAi animals, including head regression, ventral curling, epidermal lesions, and lysis (Figure 4D). WISH analysis of NTC genes *prpf3* and *prpf8* demonstrated broad parenchymal expression patterns like *prpf19*, with *prpf8* showing a noticeable stem cell expression pattern (Supplementary Figure S2). The similar phenotypes and expression patterns observed for other NTC components and factors downstream to *prpf19* suggested that the *prpf19(RNAi)* phenotype is mediated through its role in NTC, and that the NTC and spliceosome function is critical for stem cell regulation during homeostasis and regeneration.

**Figure 4.**
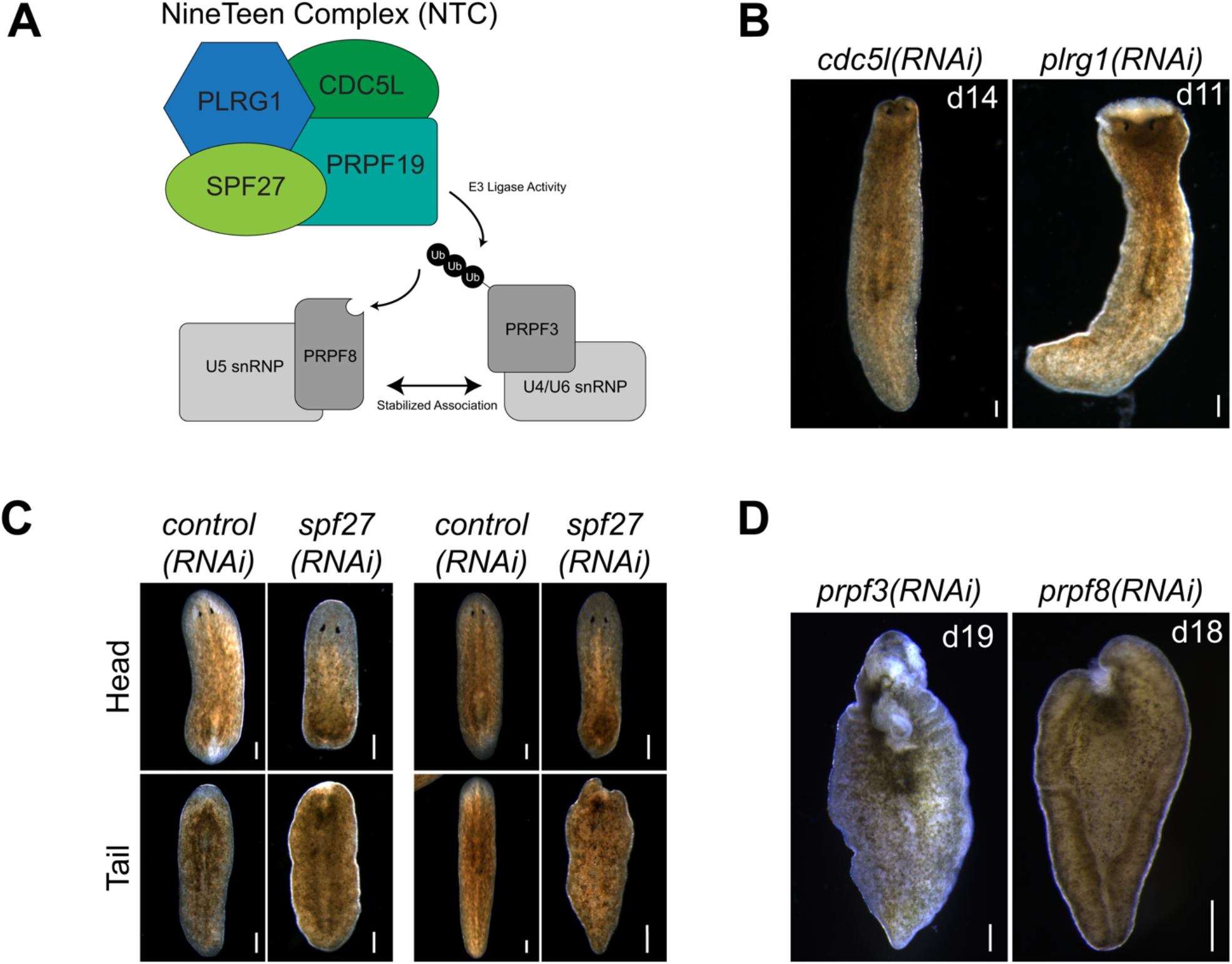
*prpf19*-associated factors and downstream targets recapitulate *prpf19(RNAi)* phenotypes. (A) Prpf19 acts as an E3 ligase in NTC, interacting with core complex members PLRG1, CDC5l, and SPF27 to modify U4/U6 snRNP subunit PRPF3 with nonproteolytic K63-linked ubiquitin chains. This ubiquityl mark stabilizes the interaction of PRPF3 with U5 snRNP subunit PRPF8 to allow the stable formation of the U4/U6.U5 tri-snRNP and the catalytic activity of the spliceosome. (B) Knockdown of indicated NTC core components *cdc5l* and *plrg1* displaying head regression (N = 19/41 and 24/44 respectively) and lysis (N = 39/41 and 20/44, respectively). (C) Knockdown of NTC core component *spf27* caused a reduced and delayed regenerative response in amputated worms. At six days post-amputation, *spf27(RNAi)* worms have smaller blastemas compared to *control(RNAi)* worms at the same time point. At day 11 post-amputation, the regenerative response in *control(RNAi)* worms is largely concluded with large blastemas and visible reformed eyespots present in trunk fragments. In comparison, *spf27(RNAi)* worms have smaller blastemas (N = 28/37 for head fragments), and tail fragments have not regenerated normal eyespots (N = 33/37). (D) Inhibition of Prpf19 target *prpf3* and ubiquityl-Prpf3 binding factor *prpf8* demonstrate phenotypes like *prpf19(RNAi)* and includes head regression (N = 19/32 and 7/32, respectively), lesions (N = 5/32 and 19/32, respectively), ventral curling (5/32 and 15/32, respectively) and lysis (N = 32/32 and 32/32, respectively). Scale bars = 200 μm.

### 3.6 Histone-modifying ubiquitin ligases are essential for regeneration and homeostasis

Ubiquitylation of histone H2B is associated with transcriptional activation and, in mammals, is mediated by the E3 ligase complex RNF20/40 (Bre1 in yeast) (Henry et al., 2003; Hwang et al., 2003). We found that planarians have a single homolog for this complex named *Smed-bre1* (referred to as *bre1* hereon). RNAi knockdown of *bre1* caused the worms to exhibit head regression and lesions prior to day 28 of treatment in 33/53 worms assayed (Figure 1B). Furthermore, most *bre1(RNAi)* worms failed to regenerate when amputated, and many lysed with 31/53 head fragments and 21/53 trunk fragments lysing by the end of the observation period (day 14 post-amputation). To investigate if *bre1(RNAi)* affected global levels of monoubiquityl-histone H2B (H2Bu1), we performed a protein blot using an H2Bub1-specific antibody. We found reduced levels of H2Bu1 in whole worm homogenates as soon as 14 days after beginning RNAi treatment (Figure 5).

**Figure 5.**
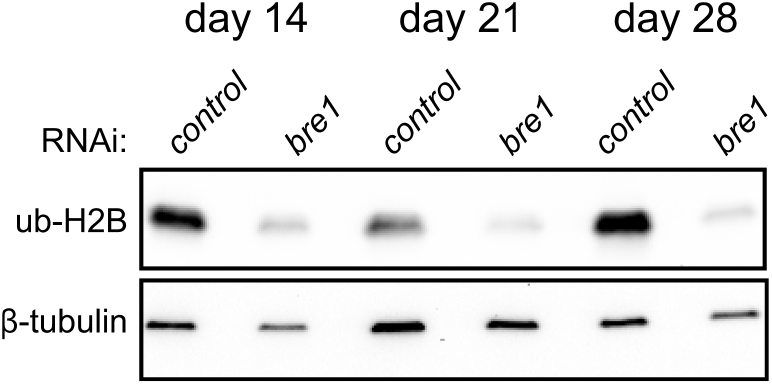
Western blot analysis shows a reduction in H2Bub1 levels following disruption of *bre1* function at days 14, 21, and 28 of RNAi treatment.

In contrast to histone H2B ubiquitylation, monoubiquitylation of histone H2A is associated with transcriptional repression. It occurs in various cellular contexts, including developmental processes, stem cell regulation, and the DNA damage response. Histone H2A is targeted for ubiquitylation by RING1 and RNF2, which act as RING E3 ligases within PRC1. PRC1 is active during development to monoubiquitylate histone H2A (Wang et al., 2004) and stably silence genes (Bunker and Kingston, 1994) (Figure 6A). We identified two candidate homologs of RING1 and RNF2 and found that depletion of each caused delayed or absent regeneration compared to controls (Figure 1C). These phenotypes were most evident in the trunk fragments where 37/58 *rnf2(RNAi)* and 19/29 *ring1(RNAi)* worms exhibited a delayed regeneration phenotype (measured by the appearance of dark eyespots) compared to 7/54 and 2/30 *control(RNAi)* worms assayed at the same regeneration time point (7 days post-amputation). Of the 37/58 *rnf2(RNAi)* trunks and 19/29 *ring1(RNAi)* trunks with regeneration defects, 13/37 and 4/19 failed to form regeneration blastemas, respectively, whereas all *control(RNAi)* worms formed normal-sized blastemas (Figure 1C). No obvious phenotypes were observed during homeostasis, even during long-term RNAi treatment (>16 feeds over eight weeks). To assess if the ubiquityl ligase activity of *rnf2* towards histone H2A was conserved in planarians, we examined bulk levels of H2Aub1 by protein blot analysis using an H2Aub1 specific antibody and observed markedly reduced levels of H2Aub1 after *rnf2* inhibition (Figure 6B). In contrast, we found that *ring1* RNAi did not appreciably affect global H2Aub1 levels (Supplementary Figure S3A), consistent with *rnf2* being the primary E3 ligase responsible for H2A ubiquitylation (de Napoles et al., 2004; Wang et al., 2004).

**Figure 6.**
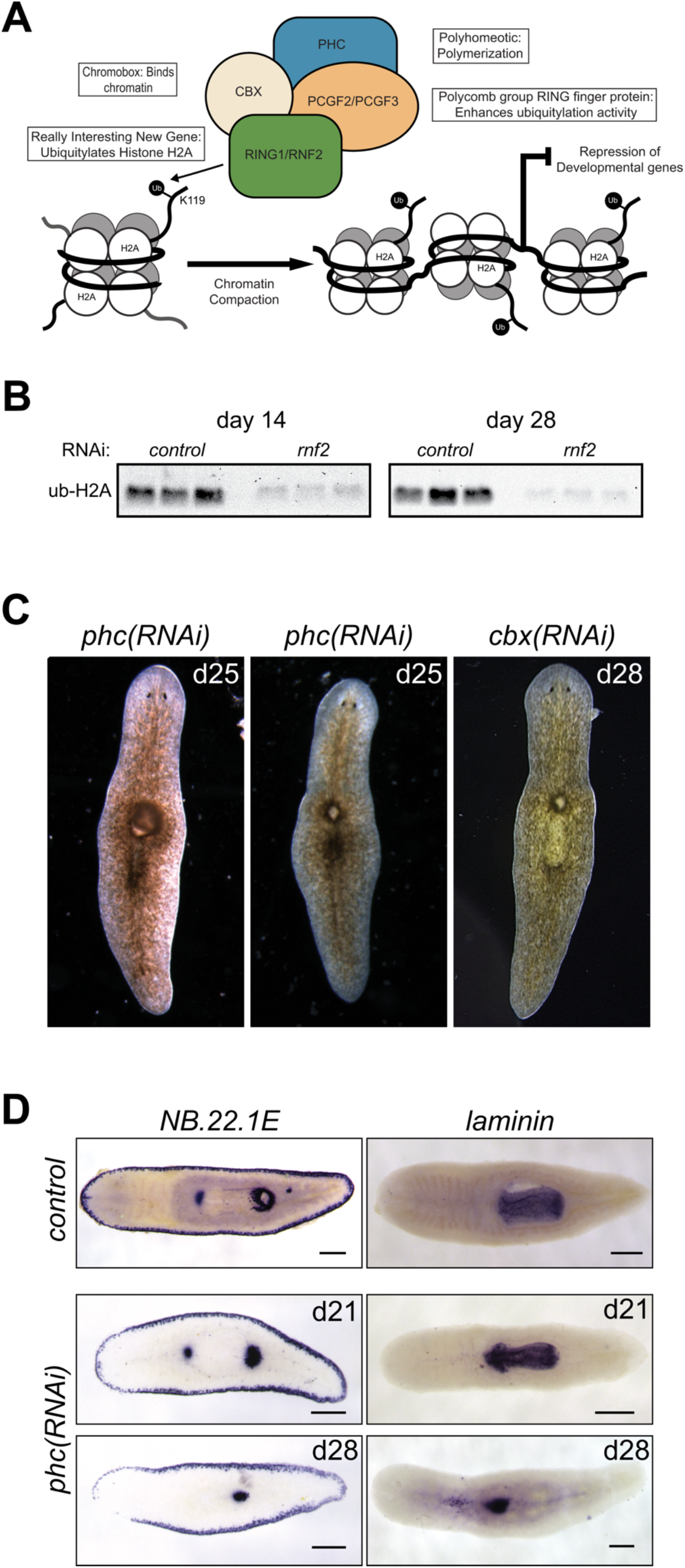
RNAi-mediated inhibition of canonical PRC1 function disrupts pharyngeal patterning and histone ubiquitylation. (A) Composition and function of PRC1. PRC1 functions to ubiquitylate histone H2A and compact chromatin to repress gene expression. (B) Western blot analysis showed a reduction in H2Aub1 levels following *rnf2* inhibition across three biological replicates and two experimental time points. (C) RNAi of PRC1 genes *phc* and *cbx* causes phenotypes of a dorsal lesion anterior to the pharynx (N = 40/95 and N = 14/32 for *phc* and *cbx*, respectively) and mislocalization of the pharynx on the dorsal surface of the worm (observed eight times in six *phc* RNAi experiments). (D) WISH to *NB*.*22*.*1E* marks the marginal adhesive gland cells, mouth opening, and a population of cells at the base of the pharynx and for *laminin*, which marks the pharynx in *control(RNAi)* animals (upper panels) and *phc(RNAi)* animals at days 21 (middle panels, N = 7-9) and 28 (bottom panels, N = 10-11) after the first RNAi treatment. Scale bars = 200 μm.

### 3.7 Disruption of canonical PRC1 subunit *phc* affects the patterning of the planarian pharyngeal body region

In vertebrates, the composition of PRC1 is variable (Gao et al., 2012); the complex is defined by which of the six mammalian paralogs of PGCF is present. PCGF2 and PCGF4 define the mammalian canonical PRC1 complex (cPRC1), which also includes one each of several chromobox (CBX) and Polyhomeotic (PHC) paralogs (Figure 6A). We identified planarian homologs for these PRC1 genes and found a single homolog each for *cbx* and *phc* and two for *pcgf* (Supplementary Table S3). To investigate if the phenotypes for *rnf2(RNAi)* and *ring1(RNAi)* were mediated through their function in cPRC1, we used RNAi to deplete *cbx, phc, pcgf2*, and *pcgf3*.

In contrast to the impaired regeneration but normal homeostasis observed after *rnf2* or *ring1* knockdown, RNAi for *phc* or *cbx* exhibited a complex homeostasis phenotype that included the abnormal appearance of a dorsal lesion anterior to the pharynx (Figure 6C). In some cases, we observed the pharynx protruding from the lesioned region and extending ectopically from the dorsal surface of the worm. As the phenotypes progressed, these RNAi worms began to exhibit defects along the body axis, showing crimped tails unable to affix to the dish and epidermal lesions. We also assayed the effect of inhibition of the canonical PRC1 genes on H2Aub1 levels. We found that inhibition of *phc* or *cbx* did not impact bulk H2Aub1 levels (Supplementary Figure S3A), suggesting that planarian cPRC1 is not responsible for bulk H2Aub1 deposition, consistent with findings in vertebrates (Fursova et al., 2019). Both *phc* and *cbx* had similar mRNA expression patterns, suggesting they have the potential to function in the same complex (Supplementary Figure S3B). This expression pattern overlapped with the diffuse parenchymal expression pattern for *rnf2* and *ring1* (Figure 2A) but had more robust expression near the planarian brain and intestinal branches, the latter of which are areas known to be enriched in stem cells.

Although similar, the penetrance of the *phc(RNAi)* phenotype was more robust than for *cbx(RNAi)*, and we chose to examine the *phc(RNAi)* phenotype further using known markers of tissue patterning. The appearance of a dorsal lesion and mislocalization of the pharynx to the dorsal surface in *phc(RNAi)* animals suggests that PRC1 may be involved in maintaining pharynx tissues or regulating genes that provide axial positioning cues to stem cell progeny during homeostatic tissue turnover. To test these hypotheses, we first examined dorsal-ventral patterning factor *bmp-4* (Gavino and Reddien, 2011) and anterior-posterior factor *ndl-3* (Rink et al., 2009) expression after disrupting *phc*. We did not observe a noticeable change in the expression pattern of these genes relative to the controls (Supplementary Figure S3C). We then further examined genes that mark specific tissues related to the pharynx, including the pharynx marker *laminin* (Adler et al., 2014) and the gene *NB*.*22*.*1E* (Tu et al., 2015), which labels marginal adhesive gland cells, the ventral mouth opening, and a population of cells near the base of the pharynx. Following *phc* inhibition, we observed that *laminin* expression was reduced to a single condensed spot of expression near the location of the dorsal lesion and a few scattered cells near the midline (Figure 6D). Likewise, we observed the specific disappearance of the *NB*.*22*.*1E*^*+*^ population of cells near the anterior end of the pharynx following *phc(RNAi)*. In contrast, expression along the body margin and ventral mouth opening was unaffected (Figure 6D). These data establish a role for PRC1 factors in maintaining specific tissue identity in a non-embryological context.

### 3.8 RNA-seq analysis after *rnf2* and *phc* RNAi inhibition reveals candidate transcriptional targets of PRC1

To investigate which genes are differentially expressed after PRC1 inhibition and to understand the transcriptional basis for the *rnf2* and *phc* RNAi phenotypes, we performed RNA-seq. We chose time points based on the phenotypic progression, quantitative PCR analysis to confirm a robust reduction in target RNAi transcript levels (not shown), and, for *rnf2(RNAi)*, protein blot analysis to ensure the RNAi treatment was reducing levels of H2Aub1. Based on these parameters, we extracted RNA after 11 days of *phc* RNAi treatment and 14 and 28 days after *rnf2* RNAi (Supplementary Figure S4A).

We identified 264 unique differentially expressed genes (126 down-regulated and 138 up-regulated) combined between the two time points sampled after *rnf2(RNAi)* (Figure 7A and 7B, Supplementary Figure S4B). Not surprisingly, an extended *rnf2* RNAi treatment led to an increase in the number of differentially expressed genes: 247 genes were differentially expressed after 28 days compared to 29 after 14 days of treatment (Figure 7A). Also, there was substantial overlap between the *rnf2(RNAi)* data sets, with 12 of 29 genes in the day 14 data set represented in the day 28 data set (Supplementary Figure S4C). After 11 days of *phc(RNAi)*, 49 genes were differentially expressed: 20 were down-regulated and 29 up-regulated (Figure 7A and 7C). Consistent with a repressive role in transcriptional regulation, more genes were significantly up-regulated when either *phc* or *rnf2* was inhibited. Importantly, *rnf2* and *phc* were each significantly down-regulated when targeted for RNAi.

**Figure 7.**
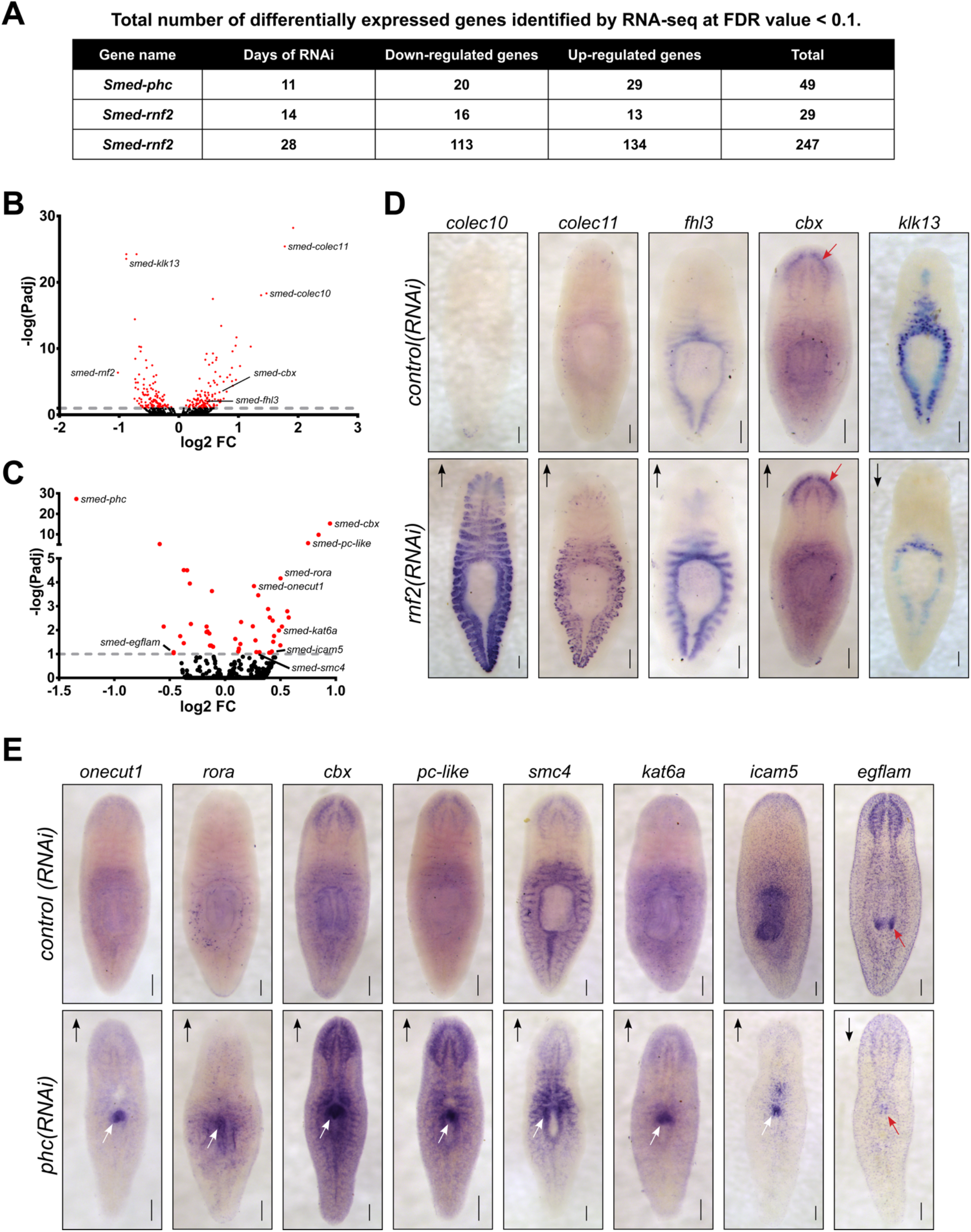
Loss of PRC1 function causes changes to gene expression levels and spatial patterning. (A) Summary of the total number of differentially expressed genes detected following RNAi knockdowns. (B) Volcano plot of differentially expressed genes after 28 days of *rnf2* RNAi treatment. (C) Volcano plot of differentially expressed genes after 28 days of *phc* RNAi. (D) WISH analysis of select genes that were differentially expressed by following *rnf2* RNAi (N = 6-12 animals per group). (E) WISH analysis of selected genes indicated to be differentially expressed after *phc* RNAi knockdown (N = 6-10 animals per group). The length of time each probe was developed was equal between target RNAi and control RNAi samples. Arrows indicate up- or down-regulated expression measured by RNA-seq. Red arrows highlight regions with changed expression after RNAi in the worm’s brain (D) and mouth (E) regions. White arrows indicate regions of ectopic gene expression after RNAi treatment. Scale bars = 200 μm.

Surprisingly, despite being predicted to function in a complex together, only a single differentially expressed gene was found in common between the *phc(RNAi)* and *rnf2(RNAi)* data sets. Intriguingly, this gene was *cbx*, which encodes a chromatin-binding element within PRC1 and was the most significantly up-regulated gene after *phc* knockdown. This overall lack of overlap between the data sets suggests that *phc* and *rnf2* regulate different processes and pathways *in vivo*, and this difference explains the disparate phenotypes observed after RNAi treatment.

To gain insight into the RNA-seq expression data, we performed Gene Ontology (GO) analysis on the differentially expressed gene set from the day 28 *rnf2(RNAi)* gene set. The down-regulated genes were significantly enriched for GO biological process terms related to metabolic and catabolic processes (Supplementary Figure S4D). Conversely, among GO terms enriched in up-regulated genes were cellular stress, especially low oxygen conditions, including “response to hypoxia” (GO:0001666), “cellular response to decreased oxygen levels” (GO:0036294), “ATF6-mediated unfolded protein response” (GO:0036500), “regulation of transcription from RNA polymerase II promoter in response to stress” (GO:0043618), “chaperone cofactor-dependent protein refolding” (GO:0051085), “protein folding in endoplasmic reticulum” (GO:0034975), and “protein refolding” (GO:0042026) (Supplementary Figure S4E). These GO annotations suggest that *rnf2* activity represses cellular responses to stress during normal homeostatic conditions and that epigenetic mechanisms facilitate the switch between homeostasis and cellular stress responses.

To investigate the spatial expression changes of differentially expressed genes from our RNA-seq data sets, we selected a subset to examine using WISH after *phc* or *rnf2* RNAi (Supplementary Table S3). For *rnf2(RNAi)*, we selected 33 differentially expressed genes that were predicted to be involved in the extracellular matrix, stress response factors, cell signaling, and chromatin regulation or transcription and assayed their expression after *rnf2* depletion. In general, *rnf2(RNAi)* caused a subtle effect on tissue-specific gene expression levels. However, in some instances, a robust change in expression occurred in *rnf2(RNAi)* worms, as seen clearly for *Smed-colec10* and *Smed-colec11*; expression of these genes is nearly undetectable in control worms as compared to *rnf2(RNAi)* worms (Figure 7D). Taken together, the GO and *in situ* analyses indicate that *rnf2* functions in broad cellular processes and that it maintains gene expression in differentiated tissues at appropriate levels.

In contrast to the mild effect on tissue-specific gene expression observed in *rnf2(RNAi)* animals, assaying mRNA expression of putative PHC target genes revealed striking changes in expression levels and spatial patterning in *phc(RNAi)* worms. We examined 11 genes using *in situ* hybridization, including genes involved in cell adhesion, cell signaling, transcription, and chromatin regulation. For 7 of these 11 genes, strong ectopic expression was observed after *phc* RNAi in the region of the worm where the dorsal lesion forms (Figure 7E). Genes ectopically expressed in this region included the cell adhesion factor *icam5*, the Cut homeobox transcription factor *onecut1*, and *roar*, which encodes an orphan nuclear receptor. We also found several chromatin regulators that were misexpressed in the region near the pharynx, including *cbx, pc-like, smc4*, and *kat6a*. Additionally, we found that the extracellular matrix protein *egflam*, which is normally expressed in the nervous system and pharynx tip, was significantly down-regulated throughout the worm. These data both validate our RNA-seq data and point to tissue-specific transcriptional changes that correlate strongly with tissue-specific functional changes.

The ectopic expression of specific factors and disruption of *NB*.*22*.*1E* and *laminin* expression at the site of tissue defects in *phc(RNAi)* worms indicates that *phc* function is required to maintain the proper specification and integrity of tissues in this body region.

## 4 Discussion

To address the role of ubiquitin signaling in stem cell regulation and regeneration in an *in vivo*, whole organism context, we performed a functional screen of the RING/U-box class of E3 ubiquitin ligases that are expressed in stem cells and progeny in *S. mediterranea*. The screen identified nine genes that demonstrated phenotypes related to stem cell function or regeneration, building on previous studies from our lab on the HECT (Henderson et al., 2015) and Cullin-RING (Strand et al., 2018) classes of E3 ligases. In addition, other studies have also uncovered roles for RING/U-box E3s in planarian regeneration, including TNF Receptor Associated Factor (TRAF)-like genes and *prpf19* (Roberts-Galbraith et al., 2016; Ziman et al., 2020).

Consistent with previous reports (Swapna et al., 2018; Ziman et al., 2020), we found that the TRAF-like family genes are expanded in planarians (Supplementary Table S1). While the evolutionary significance of this expansion remains unresolved, numerous expression and functional studies (discussed in detail in Ziman et al., 2020) in *S. mediterranea* and *Dugesia japonica* have uncovered roles for TRAF-like genes in regulating the immune response (Arnold et al., 2016; Pang et al., 2016), regeneration (Rouhana et al., 2010), and homeostasis (Ziman et al., 2020). Our finding that *traf-2A/B* is necessary for tissue homeostasis and cell survival (Table 1) agrees with Ziman et al. (2020). Further work will be necessary to resolve the mechanistic basis for how TRAF-signaling regulates homeostasis and stress and infection responses in planarians.

*prpf19* is the founding member of the large protein complex NTC. First characterized in yeast, the best-described role for NTC is in the spliceosome, where the E3 ligase function of Prpf19 is essential for forming snRNP conformations (Song et al., 2010). We found that depletion of *prpf19* caused a strong homeostasis phenotype that included head regression, lesioning, ventral curling, and lysis, all of which are morphological effects often caused by stem cell depletion (Figure 1B). We depleted other NTC member genes in this study and observed similar phenotypes to *prpf19(RNAi)*, suggesting that the *prpf19(RNAi)* phenotype is mediated through its role in NTC. Additional biochemical evidence will be necessary to demonstrate that these factors are working together formally and mediate splicing in planarian cells. In addition, we found that the stem cell population was maintained in *prpf19(RNAi)* worms, suggesting an alternative mechanism of dysregulation. This result is consistent with a previous study that observed a similar phenotype upon depletion of *prpf19*, which showed an effect on head regeneration without disrupting the stem cells (Roberts-Galbraith et al., 2016). We investigated the dynamics of the epidermal progenitor populations after *prpf19* inhibition and found the patterning of epidermal progenitor cells to be disrupted (Figure 3A).

However, further analysis of other progenitor lineages will be necessary to determine if *prpf19* has a general role in promoting differentiation or if this function is restricted to specific lineages, such as the epidermal cells. Future studies should examine the role of *prpf19* in splicing by sequencing RNAs, including pre-mRNAs, to test if there are transcripts that are especially sensitive to disruption of mRNA splicing and if mis-spliced transcripts are related to the differentiation of stem cells.

Post-transcriptional RNA processing is emerging as a major regulator of planarian stem cells and differentiation. The PIWI homolog *smedwi-2* was identified as nonessential for stem cell maintenance but necessary for proper differentiation (Reddien et al., 2005), and *smedwi-3* was shown to regulate stem cell mRNAs through two distinct activities (Kim et al., 2019). A screen of planarian ribonucleoprotein granule component homolog genes demonstrated that most were expressed in planarian stem cells and that depletion of several genes, including mRNA turnover factors, exoribonucleases, and DEAD-box RNA helicases, inhibited regeneration without affecting proliferation or stem cell maintenance (Rouhana et al., 2010). Similarly, the CCR4-NOT complex regulates the post-translational degradation of mRNAs and has been shown to have a critical role in planarian stem cell biology (Solana et al., 2013). The phenotype of CCR4-NOT complex member gene *Smed-not1* was reported to have a similar phenotype to *prpf19*, in which the animals maintained proliferative stem cells despite presenting a phenotype that suggests loss of tissue renewal (Solana et al., 2013). This study found that an additional CCR4-NOT subunit, *not4*, is critical for worm homeostasis and causes head regression and ventral curling upon inhibition (Figure 1B). This phenotype is consistent with that of *not1(RNAi)*; however, in the future, it will be necessary to examine the stem cell population using marker genes in *not4(RNAi)* worms to resolve if the phenotype is mediated through a similar mechanism. Regulation of mRNAs in planarian stem cells by several pathways, including piRNAs, deadenylation, or splicing, is crucial for homeostasis and regeneration while being dispensable for stem cell maintenance. These studies implicate post-transcriptional regulation of mRNAs in planarian stem cells as a critical process for regulating differentiation.

Epigenetic regulation of gene expression is essential during development and throughout organismal life. Our RNAi screen uncovered planarian homologs of histone-targeting RING E3 ubiquitin ligases that affected worm homeostasis and regeneration and confirmed that inhibition of *bre1* and *rnf2* reduced levels of monoubiquityl-histone H2B and H2A, respectively. This work demonstrates that activating and repressive signals provided through histone modifiers are essential for the proper specification of stem cells and maintaining cellular identity during both homeostasis and regeneration. PRC1 is a major repressive complex that works during development to ubiquitylate histone H2A, compact chromatin, and silence target gene expression (Shao et al., 1999; Francis et al., 2004; Wang et al., 2004; Pengelly et al., 2015; Tamburri et al., 2019; Blackledge et al., 2020). PRC1 function was first discovered and was best characterized as a repressor of the HOX genes during development (Lewis, 1978). The core PRC1 complex is defined by a RING and PCGF protein that forms canonical (cPRC1) or variant PRC1 (vPRC1) depending on the presence of additional factors (Conway and Bracken, 2017). The RING subunit acts as an E3 ligase that targets histone H2A, and in vertebrates is either RING1 or RNF2 (Cao et al., 2005). In contrast to *Drosophila*, we found that planarians have two homologs of RING1 and RNF2. While these are likely to be lineage-specific paralogs instead of direct homologs of each vertebrate gene, we find that, as in vertebrates, the *S. mediterranea rnf2* paralog acts as the major ligase and is responsible for the bulk of histone H2A ubiquitylation. We did not observe a noticeable difference in global H2Aub1 levels after *Smed-ring1* inhibition (Supplementary Figure S4B). However, as both genes demonstrated similar regeneration-specific phenotypes, they may share common targets or pathways.

In contrast to *rnf2(RNAi)*, when we perturbed other cPRC1 core elements *phc* and *cbx*, we did not see a reduction in bulk H2Aub1 levels by western blotting (Supplementary Figure S3A). Work in mammalian cell lines has determined that vPRC1 activity is responsible for most H2A ubiquitylation with a minimal contribution from cPRC1 complexes (Blackledge et al., 2014; Fursova et al., 2019). Invertebrates were not thought to contain vPRC1, but more recent phylogenic analysis that included a greater variety of invertebrate model organisms indicates that vPRC1 likely evolved as early as cnidarians (Gahan et al., 2020). Our protein blot results show a minimal contribution of cPRC1 genes *cbx* and *phc* to overall H2Aub1 levels *in vivo*, and the presence of two *S. mediterranea pcgf* genes support the potential existence of vPRC1 in *S. mediterranea*. Further work will be needed to elucidate the biochemical composition of PRC1, including any variant complexes that might exist, to show the direct interaction of these factors in a functional E3 ligase complex.

To gain insight into which genes are regulated by *rnf2* and *phc*, we performed RNA-seq following RNAi. Consistent with predicted roles in transcriptional repression, inhibiting the function of either gene led to more up-regulated than down-regulated differentially expressed genes. The only gene shared between the data sets was *cbx*, which was significantly up-regulated in both *rnf2* and *phc* RNAi worms. This finding suggests a possible model in which PRC1 complexes autoregulate their expression in planarians, such that the disruption of one PRC1 component causes a compensatory response involving other chromatin factors. Additionally, our GO analysis found that *rnf2* regulates genes related to the cellular stress response. When we examined the expression of differentially regulated genes from our RNA-seq data set using WISH, we observed that gene expression changes in *rnf2(RNAi)* animals occurred mainly within the endogenous expression pattern. These data support a role for RNF2 and potentially H2A ubiquitylation, in tuning transcription levels within a particular cell type, especially for pathways that are adaptive and responsive to stressful stimuli.

In contrast to *rnf2(RNAi)*, we saw dramatic shifts in the spatial expression of specific genes after *phc(RNAi)*, including several genes that showed ectopic expression near the base of the pharynx where lesions formed in *phc(RNAi)* planarians. We observed both up- and down-regulation of genes that encode extracellular matrix and intercellular adhesion molecules such as *intercellular adhesion molecule 5* and *pikachurin*, respectively, suggesting that their dysregulation is likely linked to the formation of the dorsal lesion seen. Interestingly, RNA-seq did not detect differential expression of *foxA*, which is required to specify pharyngeal tissues, nor did we find any overlap with our data set and factors that are upregulated after the amputation of the pharynx (Adler et al., 2014). The anatomical location of misexpressed genes after *phc* RNAi correlates strongly with the location of pharynx progenitors, but our RNA-seq data set did not recover differential expression of known pharynx genes. It is possible that the pharynx specification gene response is temporally shifted in *phc(RNAi)* animals, which could be tested by increasing the number of time points. In addition, assaying the direct localization of PHC in the genome could uncover loci that are regulated by PHC.

The ectopically expressed genes also included regulators of cellular specification, including nuclear receptors, transcription factors, and chromatin modifiers. One gene we identified as being misexpressed after *phc* depletion was that encoding the nuclear factor *onecut1*, a CUT, and homeobox domain-containing transcription factor that promotes hepatocyte proliferation, remodels chromatin accessibility, and promotes tumor growth in colorectal cancers (Jiang et al., 2019; van der Raadt et al., 2019; Peng et al., 2020). Based on its role in regulating transcription and tissue identity in other animal models, we suspect it may be contributing to the change in patterning near the pharynx. Future investigation will elucidate if *onecut1* misexpression drives regional tissue misspecification and if inhibition of *onecut1* suppresses the *phc(RNAi)* phenotype.

PRC1 regulates gene expression through the post-translational ubiquitylation of histone H2A and by compacting chromatin. A deeper understanding of the functional role of this complex in regulating regeneration will require uncovering its specific biochemical composition and genomic elements it targets. In the future, assays like ChIP-seq (Lee et al., 2006) or CUT&RUN (Skene and Henikoff, 2017) could be used to identify the localization of H2Aub1 in the planarian genome and inform how genes are regulated epigenetically to promote a robust regenerative response during injury and tissue re-specification and remodeling.

## 6 Author Contributions

J.M.A. and R.M.Z. conceptualized the study and designed experiments. J.M.A., M.B., E.B., C.C.M., S.A.Z., and I.I. conducted experiments. J.M.A. and R.M.Z. wrote the original draft of the manuscript. The manuscript was reviewed and edited by J.M.A., E.M.D., and R.M.Z. Funding for the project was acquired by R.M.Z.

## 8 Acknowledgments

We thank Celeste Romero for help with RNAi experiments, Kelly Ross for helpful discussions, Mohammad Auwal and Mallory Palatucci for comments on the manuscript, and Catherine May from Boston Children’s Hospital for helping to set up the RNA-seq analysis and providing template R scripts.

## 10 Supplementary Material

Supplementary Material includes four files: a single .pdf file containing four supplementary figures (Supplementary Figures S1-S4) and three supplementary tables in .xls format (Supplementary Tables S1-S3).

## 11 Data Availability Statement

The datasets generated and analyzed for this study can be found in the NCBI database under BioProject PRJNA768725.

## References

Abnave, P., Aboukhatwa, E., Kosaka, N., Thompson, J., Hill, M.A., and Aboobaker, A.A. (2017). Epithelial-mesenchymal transition transcription factors control pluripotent adult stem cell migration in vivo in planarians. Development 144(19), 3440–3453. doi: 10.1242/dev.154971.

Adler, C.E., and Sánchez Alvarado, A. (2018). Systemic RNA Interference in Planarians by Feeding of dsRNA Containing Bacteria. Methods Mol Biol 1774, 445–454. doi: 10.1007/978-1-4939-7802-1_17.

Adler, C.E., Seidel, C.W., McKinney, S.A., and Sánchez Alvarado, A. (2014). Selective amputation of the pharynx identifies a FoxA-dependent regeneration program in planaria. Elife 3, e02238. doi: 10.7554/eLife.02238.

Aravind, L., and Koonin, E.V. (2000). The U box is a modified RING finger - a common domain in ubiquitination. Curr Biol 10(4), 132–134. doi: http://dx.doi.org/10.1016/S0960-9822(00)00398-5.

Arnold, C.P., Merryman, M.S., Harris-Arnold, A., McKinney, S.A., Seidel, C.W., Loethen, S., et al. (2016). Pathogenic shifts in endogenous microbiota impede tissue regeneration via distinct activation of TAK1/MKK/p38. Elife 5. doi: 10.7554/eLife.16793.

Baguñà, J., Saló, E., and Auladell, C. (1989). Regeneration and pattern formation in planarians. III. Evidence that neoblasts are totipotent stem cells and the source of blastema cells. Development 107, 77–86.

Blackledge, N.P., Farcas, A.M., Kondo, T., King, H.W., McGouran, J.F., Hanssen, L.L.P., et al. (2014). Variant PRC1 complex-dependent H2A ubiquitylation drives PRC2 recruitment and polycomb domain formation. Cell 157(6), 1445–1459. doi: 10.1016/j.cell.2014.05.004.

Blackledge, N.P., Fursova, N.A., Kelley, J.R., Huseyin, M.K., Feldmann, A., and Klose, R.J. (2020). PRC1 Catalytic Activity Is Central to Polycomb System Function. Mol Cell 77(4), 857–874 e859. doi: 10.1016/j.molcel.2019.12.001.

Blum, M., Chang, H.Y., Chuguransky, S., Grego, T., Kandasaamy, S., Mitchell, A., et al. (2021). The InterPro protein families and domains database: 20 years on. Nucleic Acids Res 49(D1), D344-D354. doi: 10.1093/nar/gkaa977.

Boser, A., Drexler, H.C., Reuter, H., Schmitz, H., Wu, G., Scholer, H.R., et al. (2013). SILAC proteomics of planarians identifies Ncoa5 as a conserved component of pluripotent stem cells. Cell Rep 5(4), 1142–1155. doi: 10.1016/j.celrep.2013.10.035.

Brandl, H., Moon, H., Vila-Farre, M., Liu, S.Y., Henry, I., and Rink, J.C. (2016). PlanMine--a mineable resource of planarian biology and biodiversity. Nucleic Acids Res 44(D1), D764-773. doi: 10.1093/nar/gkv1148.

Bray, N.L., Pimentel, H., Melsted, P., and Pachter, L. (2016). Near-optimal probabilistic RNA-seq quantification. Nat Biotechnol 34(5), 525–527. doi: 10.1038/nbt.3519.

Brown, D.D.R., and Pearson, B.J. (2015). “One FISH, dFISH, Three FISH: Sensitive Methods of Whole-Mount Fluorescent In Situ Hybridization in Freshwater Planarians,” in In Situ Hybridization Methods.), 127–150.

Bunker, C.A., and Kingston, R.E. (1994). Transcriptional repression by Drosophila and mammalian Polycomb group proteins in transfected mammalian cells. Mol Cell Biol 14(3), 1721–1732. doi: 10.1128/mcb.14.3.1721-1732.1994.

Cao, R., Tsukada, Y., and Zhang, Y. (2005). Role of Bmi-1 and Ring1A in H2A ubiquitylation and Hox gene silencing. Mol Cell 20(6), 845–854. doi: 10.1016/j.molcel.2005.12.002.

Cebrià, F., and Newmark, P.A. (2005). Planarian homologs of netrin and netrin receptor are required for proper regeneration of the central nervous system and the maintenance of nervous system architecture. Development 132(16), 3691–3703. doi: 10.1242/dev.01941.

Ciechanover, A., Finley, D., and Varshavsky, A. (1984). Ubiquitin dependence of selective protein degradation demonstrated in the mammalian cell cycle mutant ts85. Cell 37(1), 57–66. doi: 10.1016/0092-8674(84)90300-3.

Conway, E.M., and Bracken, A.P. (2017). “Unraveling the Roles of Canonical and Noncanonical PRC1 Complexes,” in Polycomb Group Proteins.), 57–80.

David, Y., Ternette, N., Edelmann, M.J., Ziv, T., Gayer, B., Sertchook, R., et al. (2011). E3 ligases determine ubiquitination site and conjugate type by enforcing specificity on E2 enzymes. J Biol Chem 286(51), 44104–44115. doi: 10.1074/jbc.M111.234559.

de Napoles, M., Mermoud, J.E., Wakao, R., Tang, Y.A., Endoh, M., Appanah, R., et al. (2004). Polycomb group proteins Ring1A/B link ubiquitylation of histone H2A to heritable gene silencing and X inactivation. Dev Cell 7(5), 663–676. doi: 10.1016/j.devcel.2004.10.005.

Duncan, E.M., Chitsazan, A.D., Seidel, C.W., and Sánchez Alvarado, A. (2015). Set1 and MLL1/2 Target Distinct Sets of Functionally Different Genomic Loci In Vivo. Cell Rep 13(12), 2741–2755. doi: 10.1016/j.celrep.2015.11.059.

Eisenhoffer, G.T., Kang, H., and Sánchez Alvarado, A. (2008). Molecular analysis of stem cells and their descendants during cell turnover and regeneration in the planarian Schmidtea mediterranea. Cell Stem Cell 3(3), 327–339. doi: 10.1016/j.stem.2008.07.002.

Elliott, S.A., and Sánchez Alvarado, A. (2013). The history and enduring contributions of planarians to the study of animal regeneration. Wiley Interdiscip Rev Dev Biol 2(3), 301–326. doi: 10.1002/wdev.82.

Endoh, M., Endo, T.A., Endoh, T., Isono, K., Sharif, J., Ohara, O., et al. (2012). Histone H2A mono-ubiquitination is a crucial step to mediate PRC1-dependent repression of developmental genes to maintain ES cell identity. PLoS Genet 8(7), e1002774. doi: 10.1371/journal.pgen.1002774.

Fernandez-Taboada, E., Rodriguez-Esteban, G., Salo, E., and Abril, J.F. (2011). A proteomics approach to decipher the molecular nature of planarian stem cells. BMC Genomics 12, 133. doi: 10.1186/1471-2164-12-133.

Fincher, C.T., Wurtzel, O., de Hoog, T., Kravarik, K.M., and Reddien, P.W. (2018). Cell type transcriptome atlas for the planarian Schmidtea mediterranea. Science 360(6391). doi: 10.1126/science.aaq1736.

Francis, N.J., Kingston, R.E., and Woodcock, C.L. (2004). Chromatin compaction by a polycomb group protein complex. Science 306(5701), 1574–1577. doi: 10.1126/science.1100576.

Fursova, N.A., Blackledge, N.P., Nakayama, M., Ito, S., Koseki, Y., Farcas, A.M., et al. (2019). Synergy between Variant PRC1 Complexes Defines Polycomb-Mediated Gene Repression. Mol Cell 74(5), 1020–1036 e1028. doi: 10.1016/j.molcel.2019.03.024.

Gahan, J.M., Rentzsch, F., and Schnitzler, C.E. (2020). The genetic basis for PRC1 complex diversity emerged early in animal evolution. Proc Natl Acad Sci U S A. doi: 10.1073/pnas.2005136117.

Gao, Z., Zhang, J., Bonasio, R., Strino, F., Sawai, A., Parisi, F., et al. (2012). PCGF homologs, CBX proteins, and RYBP define functionally distinct PRC1 family complexes. Mol Cell 45(3), 344–356. doi: 10.1016/j.molcel.2012.01.002.

Gavino, M.A., and Reddien, P.W. (2011). A Bmp/Admp regulatory circuit controls maintenance and regeneration of dorsal-ventral polarity in planarians. Curr Biol 21(4), 294–299. doi: 10.1016/j.cub.2011.01.017.

Gurley, K.A., Rink, J.C., and Sánchez Alvarado, A. (2008). β-Catenin Defines Head Versus Tail Identity During Planarian Regeneration and Homeostasis. Science 319(5861), 323–327.

Henderson, J.M., Nisperos, S.V., Weeks, J., Ghulam, M., Marin, I., and Zayas, R.M. (2015). Identification of HECT E3 ubiquitin ligase family genes involved in stem cell regulation and regeneration in planarians. Dev Biol 404(2), 21–34. doi: 10.1016/j.ydbio.2015.04.021.

Henry, K.W., Wyce, A., Lo, W.S., Duggan, L.J., Emre, N.C., Kao, C.F., et al. (2003). Transcriptional activation via sequential histone H2B ubiquitylation and deubiquitylation, mediated by SAGA-associated Ubp8. Genes Dev 17(21), 2648–2663. doi: 10.1101/gad.1144003.

Higgins, R., Gendron, J.M., Rising, L., Mak, R., Webb, K., Kaiser, S.E., et al. (2015). The Unfolded Protein Response Triggers Site-Specific Regulatory Ubiquitylation of 40S Ribosomal Proteins. Mol Cell 59(1), 35–49. doi: 10.1016/j.molcel.2015.04.026.

Huber, W., Carey, V.J., Gentleman, R., Anders, S., Carlson, M., Carvalho, B.S., et al. (2015). Orchestrating high-throughput genomic analysis with Bioconductor. Nat Methods 12(2), 115–121. doi: 10.1038/nmeth.3252.

Hwang, W.W., Venkatasubrahmanyam, S., Ianculescu, A.G., Tong, A., Boone, C., and Madhani, H.D. (2003). A Conserved RING Finger Protein Required for Histone H2B Monoubiquitination and Cell Size Control. Molecular Cell 11(1), 261–266. doi: 10.1016/s1097-2765(02)00826-2.

Ivankovic, M., Haneckova, R., Thommen, A., Grohme, M.A., Vila-Farré, M., Werner, S., et al. (2019). Model systems for regeneration: planarians. Development 146(17), dev167684. doi: 10.1242/dev.167684.

Jiang, K., Jiao, Y., Liu, Y., Fu, D., Geng, H., Chen, L., et al. (2019). HNF6 promotes tumor growth in colorectal cancer and enhances liver metastasis in mouse model. J Cell Physiol 234(4), 3675–3684. doi: 10.1002/jcp.27140.

Kim, I.V., Duncan, E.M., Ross, E.J., Gorbovytska, V., Nowotarski, S.H., Elliott, S.A., et al. (2019). Planarians recruit piRNAs for mRNA turnover in adult stem cells. Genes Dev 33(21-22), 1575-1590. doi: 10.1101/gad.322776.118.

King, R.S., and Newmark, P.A. (2013). In situ hybridization protocol for enhanced detection of gene expression in the planarian Schmidtea mediterranea. BMC Developmental Biology 13(8).

Labbé, R.M., Irimia, M., Currie, K.W., Lin, A., Zhu, S.J., Brown, D.D., et al. (2012). A comparative transcriptomic analysis reveals conserved features of stem cell pluripotency in planarians and mammals. Stem Cells 30(8), 1734–1745. doi: 10.1002/stem.1144.

Lauter, G., Soll, I., and Hauptmann, G. (2011). Two-color fluorescent in situ hybridization in the embryonic zebrafish brain using differential detection systems. BMC Dev Biol 11, 43. doi: 10.1186/1471-213X-11-43.

Lee, T.I., Johnstone, S.E., and Young, R.A. (2006). Chromatin immunoprecipitation and microarray-based analysis of protein location. Nat Protoc 1(2), 729–748. doi: 10.1038/nprot.2006.98.

Lewis, E.B. (1978). A gene complex controlling segmentation in Drosophila. Nature 276(5688), 565–570. doi: 10.1038/276565a0.

Li, W., Bengtson, M.H., Ulbrich, A., Matsuda, A., Reddy, V.A., Orth, A., et al. (2008). Genome-wide and functional annotation of human E3 ubiquitin ligases identifies MULAN, a mitochondrial E3 that regulates the organelle’s dynamics and signaling. PLoS One 3(1), e1487. doi: 10.1371/journal.pone.0001487.

Liu, S.Y., Selck, C., Friedrich, B., Lutz, R., Vila-Farré, M., Dahl, A., et al. (2013). Reactivating head regrowth in a regeneration-deficient planarian species. Nature 500(7460), 81–84. doi: 10.1038/nature12414.

Livak, K.J., and Schmittgen, T.D. (2001). Analysis of relative gene expression data using real-time quantitative PCR and the 2(-Delta Delta C(T)) Method. Methods 25(4), 402–408. doi: 10.1006/meth.2001.1262.

Lorick, K.L., Jensen, J.P., Fang, S., Ong, A.M., Hatakeyama, S., and Weissman, A.M. (1999). RING fingers mediate ubiquitin-conjugating enzyme (E2)-dependent ubiquitination. Proc Natl Acad Sci U S A 96(20), 11364–11369. doi: 10.1073/pnas.96.20.11364.

Love, M.I., Huber, W., and Anders, S. (2014). Moderated estimation of fold change and dispersion for RNA-seq data with DESeq2. Genome Biol 15(12), 550. doi: 10.1186/s13059-014-0550-8.

Lu, X., and Legerski, R.J. (2007). The Prp19/Pso4 core complex undergoes ubiquitylation and structural alterations in response to DNA damage. Biochem Biophys Res Commun 354(4), 968–974. doi: 10.1016/j.bbrc.2007.01.097.

Merryman, M.S., Sánchez Alvarado, A., and Jenkin, J.C. (2018). Culturing Planarians in the Laboratory. Methods Mol Biol 1774, 241–258. doi: 10.1007/978-1-4939-7802-1_5.

Nakayama, K.I., and Nakayama, K. (2005). Regulation of the cell cycle by SCF-type ubiquitin ligases. Semin Cell Dev Biol 16(3), 323–333. doi: 10.1016/j.semcdb.2005.02.010.

Onal, P., Grun, D., Adamidi, C., Rybak, A., Solana, J., Mastrobuoni, G., et al. (2012). Gene expression of pluripotency determinants is conserved between mammalian and planarian stem cells. EMBO J 31(12), 2755–2769. doi: 10.1038/emboj.2012.110.

Pang, Q., Gao, L., Hu, W., An, Y., Deng, H., Zhang, Y., et al. (2016). De Novo Transcriptome Analysis Provides Insights into Immune Related Genes and the RIG-I-Like Receptor Signaling Pathway in the Freshwater Planarian (Dugesia japonica). PLoS One 11(3), e0151597. doi: 10.1371/journal.pone.0151597.

Pearson, B.J., Eisenhoffer, G.T., Gurley, K.A., Rink, J.C., Miller, D.E., and Sánchez Alvarado, A. (2009). Formaldehyde-based whole-mount in situ hybridization method for planarians. Dev Dyn 238(2), 443–450. doi: 10.1002/dvdy.21849.

Pellettieri, J., Fitzgerald, P., Watanabe, S., Mancuso, J., Green, D.R., and Sánchez Alvarado, A. (2010). Cell death and tissue remodeling in planarian regeneration. Dev Biol 338(1), 76–85. doi: 10.1016/j.ydbio.2009.09.015.

Peng, Y.C., Lv, T.H., Du, Z.K., Cun, X.N., and Yang, K.M. (2020). Liver Macrophages Stimulate the Expression of Hepatocyte Nuclear Factor-6 and Promote Hepatocyte Proliferation at the Early Stage of Liver Regeneration. Bull Exp Biol Med 170(1), 40–45. doi: 10.1007/s10517-020-05000-7.

Pengelly, A.R., Kalb, R., Finkl, K., and Muller, J. (2015). Transcriptional repression by PRC1 in the absence of H2A monoubiquitylation. Genes Dev 29(14), 1487–1492. doi: 10.1101/gad.265439.115.

Plass, M., Solana, J., Wolf, F.A., Ayoub, S., Misios, A., Glarzar, P., et al. (2018). Cell type atlas and lineage tree of a whole complex animal by single-cell transcriptomics. Science. doi: 10.1126/science.aaq1723(2018).

Reddien, P.W. (2018). The Cellular and Molecular Basis for Planarian Regeneration. Cell 175(2), 327–345. doi: 10.1016/j.cell.2018.09.021.

Reddien, P.W., Oviedo, N.J., Jennings, J.R., Jenkin, J.C., and Sánchez Alvarado, A. (2005). SMEDWI-2 is a PIWI-like protein that regulates planarian stem cells. Science 310(5752), 1327–1330. doi: 10.1126/science.1116110.

Rink, J.C., Gurley, K.A., Elliott, S.A., and Sánchez Alvarado, A. (2009). Planarian Hh signaling regulates regeneration polarity and links Hh pathway evolution to cilia. Science 326(5958), 1406–1410. doi: 10.1126/science.1178712.

Roberts-Galbraith, R.H., Brubacher, J.L., and Newmark, P.A. (2016). A functional genomics screen in planarians reveals regulators of whole-brain regeneration. Elife 5. doi: 10.7554/eLife.17002.

Rouhana, L., Shibata, N., Nishimura, O., and Agata, K. (2010). Different requirements for conserved post-transcriptional regulators in planarian regeneration and stem cell maintenance. Dev Biol 341(2), 429–443. doi: 10.1016/j.ydbio.2010.02.037.

Rouhana, L., Weiss, J.A., Forsthoefel, D.J., Lee, H., King, R.S., Inoue, T., et al. (2013). RNA interference by feeding in vitro-synthesized double-stranded RNA to planarians: methodology and dynamics. Dev Dyn 242(6), 718–730. doi: 10.1002/dvdy.23950.

Rozanski, A., Moon, H., Brandl, H., Martin-Duran, J.M., Grohme, M.A., Huttner, K., et al. (2019). PlanMine 3.0-improvements to a mineable resource of flatworm biology and biodiversity. Nucleic Acids Res 47(D1), D812–D820. doi: 10.1093/nar/gky1070.

Saló, E., and Baguñà, J. (1985). Cell movement in intact and regenerating planarians. Quantitation using chromosomal, nuclear and cytoplasmic markers. Development 89(1), 57–70. doi: 10.1242/dev.89.1.57.

Shao, Z., Raible, F., Mollaaghababa, R., Guyon, J.R., Wu, C.-t., Bender, W., et al. (1999). Stabilization of Chromatin Structure by PRC1, a Polycomb Complex. Cell 98(1), 37–46. doi: 10.1016/s0092-8674(00)80604-2.

Simoes, A.E., Pereira, D.M., Amaral, J.D., Nunes, A.F., Gomes, S.E., Rodrigues, P.M., et al. (2013). Efficient recovery of proteins from multiple source samples after TRIzol((R)) or TRIzol((R))LS RNA extraction and long-term storage. BMC Genomics 14, 181. doi: 10.1186/1471-2164-14-181.

Skene, P.J., and Henikoff, S. (2017). An efficient targeted nuclease strategy for high-resolution mapping of DNA binding sites. Elife 6. doi: 10.7554/eLife.21856.

Solana, J., Gamberi, C., Mihaylova, Y., Grosswendt, S., Chen, C., Lasko, P., et al. (2013). The CCR4-NOT complex mediates deadenylation and degradation of stem cell mRNAs and promotes planarian stem cell differentiation. PLoS Genet 9(12), e1004003. doi: 10.1371/journal.pgen.1004003.

Song, E.J., Werner, S.L., Neubauer, J., Stegmeier, F., Aspden, J., Rio, D., et al. (2010). The Prp19 complex and the Usp4Sart3 deubiquitinating enzyme control reversible ubiquitination at the spliceosome. Genes Dev 24(13), 1434–1447. doi: 10.1101/gad.1925010.

Strand, N.S., Allen, J.M., Ghulam, M., Taylor, M.R., Munday, R.K., Carrillo, M., et al. (2018). Dissecting the function of Cullin-RING ubiquitin ligase complex genes in planarian regeneration. Dev Biol 433(2), 210–217. doi: 10.1016/j.ydbio.2017.10.011.

Swapna, L.S., Molinaro, A.M., Lindsay-Mosher, N., Pearson, B.J., and Parkinson, J. (2018). Comparative transcriptomic analyses and single-cell RNA sequencing of the freshwater planarian Schmidtea mediterranea identify major cell types and pathway conservation. Genome Biol 19(1), 124. doi: 10.1186/s13059-018-1498-x.

Tamburri, S., Lavarone, E., Fernandez-Perez, D., Conway, E., Zanotti, M., Manganaro, D., et al. (2019). Histone H2AK119 Mono-Ubiquitination Is Essential for Polycomb-Mediated Transcriptional Repression. Mol Cell. doi: 10.1016/j.molcel.2019.11.021.

Tu, K.C., Cheng, L.C., h, T.K.V., Lange, J.J., McKinney, S.A., Seidel, C.W., et al. (2015). Egr-5 is a post-mitotic regulator of planarian epidermal differentiation. Elife 4, e10501. doi: 10.7554/eLife.10501.

van der Raadt, J., van Gestel, S.H.C., Nadif Kasri, N., and Albers, C.A. (2019). ONECUT transcription factors induce neuronal characteristics and remodel chromatin accessibility. Nucleic Acids Res 47(11), 5587–5602. doi: 10.1093/nar/gkz273.

van Wolfswinkel, J.C., Wagner, D.E., and Reddien, P.W. (2014). Single-cell analysis reveals functionally distinct classes within the planarian stem cell compartment. Cell Stem Cell 15(3), 326–339. doi: 10.1016/j.stem.2014.06.007.

Wagner, D.E., Wang, I.E., and Reddien, P.W. (2011). Clonogenic neoblasts are pluripotent adult stem cells that underlie planarian regeneration. Science 332(6031), 811–816. doi: 10.1126/science.1203983.

Wang, H., Wang, L., Erdjument-Bromage, H., Vidal, M., Tempst, P., Jones, R.S., et al. (2004). Role of histone H2A ubiquitination in Polycomb silencing. Nature 431(7010), 873–878. doi: 10.1038/nature02985.

Werner, A., Manford, A.G., and Rape, M. (2017). Ubiquitin-Dependent Regulation of Stem Cell Biology. Trends Cell Biol 27(8), 568–579. doi: 10.1016/j.tcb.2017.04.002.

Zayas, R.M., Hernandez, A., Habermann, B., Wang, Y., Stary, J.M., and Newmark, P.A. (2005). The planarian Schmidtea mediterranea as a model for epigenetic germ cell specification: analysis of ESTs from the hermaphroditic strain. Proc Natl Acad Sci U S A 102(51), 18491–18496. doi: 10.1073/pnas.0509507102.

Ziman, B., Barghouth, P.G., Maciel, E.I., and Oviedo, N.J. (2020). TRAF-like Proteins Regulate Cellular Survival in the Planarian Schmidtea mediterranea. iScience 23(11), 101665. doi: 10.1016/j.isci.2020.101665.

