## Supplementary Figures S1-4 for "RNAi screen of RING/U-box domain ubiquitin ligases identifies critical regulators of tissue regeneration in planarians"

### *Supplementary Material*

#### 1 Supplemental Figure Legends

**Supplemental Figure S1.** (A-B) Double fluorescence *in situ* hybridization showing co-expression of *prpf19* (magenta) with marker genes (green) for stem cells (*piwi-1*, *h2b*), early epidermal progeny marker (*prog-1*), and late epidermal progeny marker (*agat-1*). (C) Western blot analysis using an anti-ubiquitin antibody, which detects all forms of ubiquitin and ubiquitylated proteins. Whole worm homogenate protein extracts were obtained from *control(RNAi)* or *prp19f(RNAi)* animals at days indicated. Scale bars in A-B = 20 µm. (D) RT-qPCR experiments measuring the relative expression of epidermal lineage markers, *zfp-1*, *prog-1*, *agat-1*, or *vim-1* in *control(RNAi)* or prpf19(RNAi) planarians following 14 days of RNAi treatment. Graph shows mean ± SD expression levels relative to the controls. *p-value < 0.05, ***p-value < 0.0001, Student's t-test with Holm-Sidak correction for multiple comparisons.

**Supplemental Figure S2.** WISH to NTC core elements *cdc5l*, *pflg1*, *spf27*, and spliceosomal RNP members *prpf3* and *prpf8*. Scale bars = 200 µm.

**Supplemental Figure S3.** (A) Western blot analysis shows no reduction in H2Aub1 levels following disruption of canonical PRC1 genes *ring1*, *cbx*, and *phc* after 21 days of RNAi treatment. (B) WISH analysis showing expression patterns for PRC1 genes *phc* and *cbx*. (C) WISH marker gene analysis for dorsoventral marker *bmp4* and anterior-posterior marker *ndl-3* showed no change in expression domains following *phc* RNAi. Scale bars = 200 µm.

**Supplemental Figure S4.** (A) Feeding and sampling schedule for RNA-seq experiments. Worms were fed twice a week and sampled for RNA extraction at the days indicated by the red arrowheads. (B) Volcano plot of differentially expressed genes after 28 days of *rnf2* RNAi treatment. (C) Venn diagram showing the overlap between day 28 and day 14 sample sets for *rnf2(RNAi)*. (D) GO analysis of down-regulated genes in *rnf2(RNAi)* worms after 28 days of treatment. (E) GO analysis of up-regulated genes in *rnf2(RNAi)* worms after 28 days of treatment.

**Supplementary Figure S1**

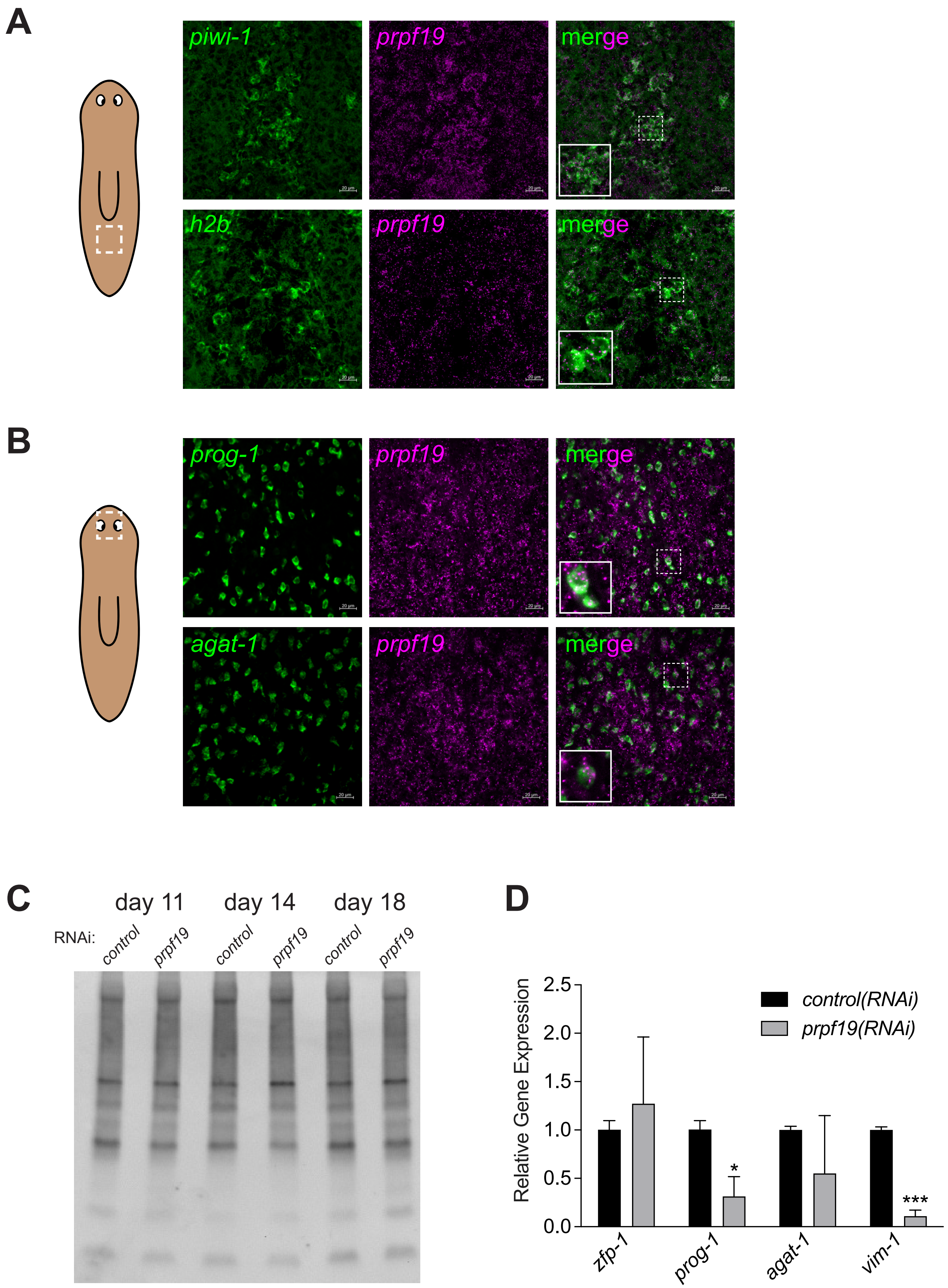

**Supplementary Figure S2**

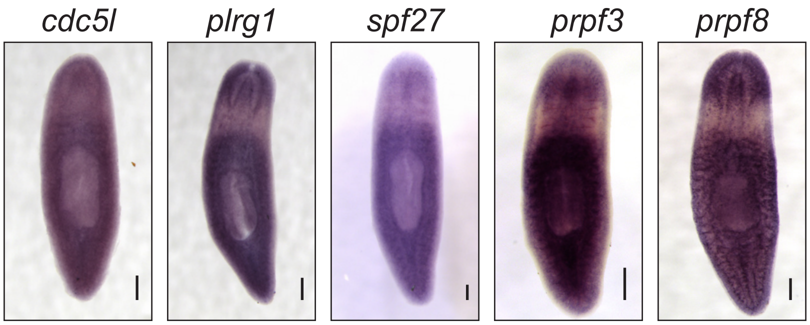

**Supplementary Figure S3**

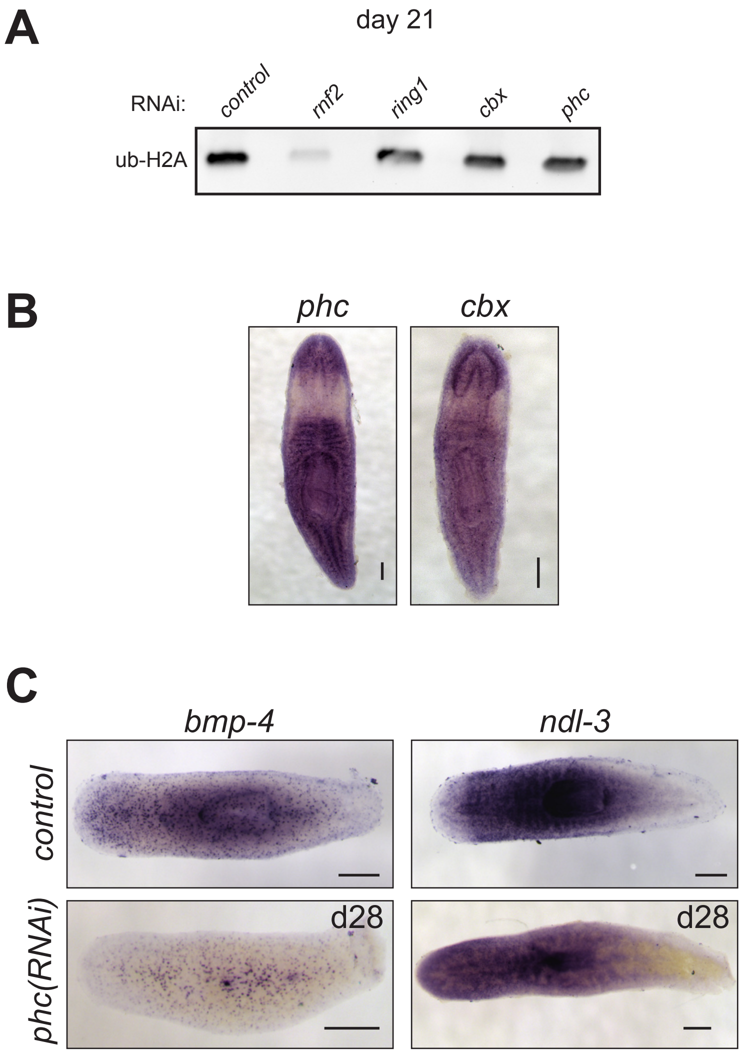

### Supplementary Figure S4

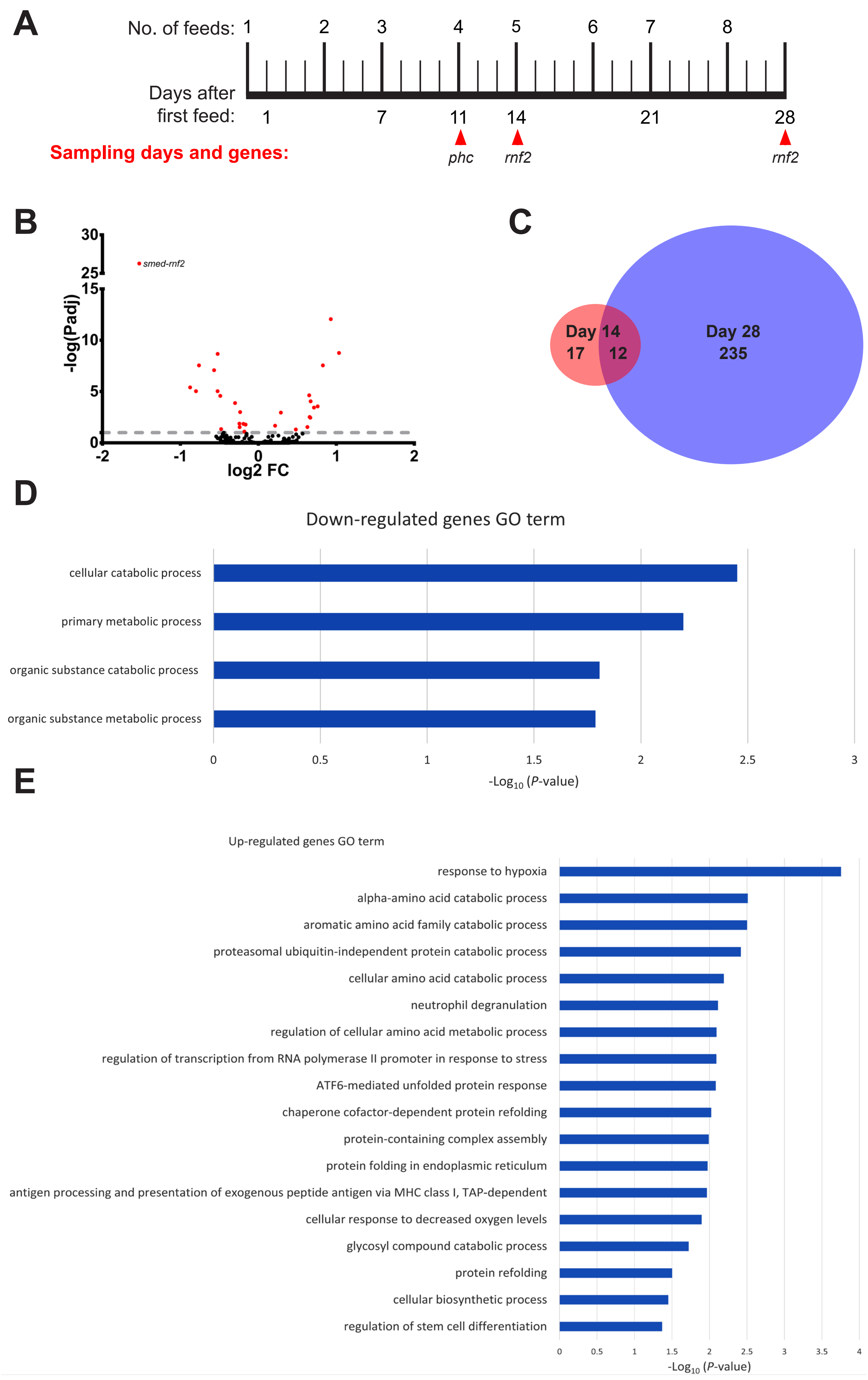
